# Cell-type-specific patterns and consequences of somatic mutation in development and aging brain

**DOI:** 10.1101/2025.05.30.656844

**Authors:** Andrea J. Kriz, Shulin Mao, Diane D. Shao, Daniel A. Snellings, Rebecca E. Andersen, Guanlan Dong, Chanthia C. Ma, Hayley E. Cline, August Yue Huang, Eunjung Alice Lee, Christopher A. Walsh

**Affiliations:** Division of Genetics and Genomics, Boston Children’s Hospital, Harvard Medical School, Boston, MA 02115, USA; Howard Hughes Medical Institute, Boston Children’s Hospital, Harvard Medical School, Boston, MA 02115, USA; Manton Center for Orphan Disease Research, Boston Children’s Hospital, Boston, MA 02115, USA; Department of Pediatrics, Harvard Medical School, Boston, MA 02115, USA; Program in Biological and Biomedical Sciences, Harvard Medical School, Boston, MA 02115, USA; Department of Neurology, Boston Children’s Hospital, Boston, MA 02115, USA; Bioinformatics and Integrative Genomics Program, Harvard Medical School; Boston, MA 02115, USA; Broad Institute of MIT and Harvard, Cambridge, MA 02142, USA; Harvard-MIT MD/PhD program, Harvard Medical School, Boston, MA

## Abstract

Elucidating the role of somatic mutations in cancer, healthy tissues, and aging depends on methods that can accurately characterize somatic mosaicism across different cell types, as well as assay their impact on cellular function. Current technologies to study cell-type-specific somatic mutations within tissues are low-throughput. We developed Duplex-Multiome, incorporating duplex consensus sequencing to accurately identify somatic single-nucleotide variants (sSNV) from the same nucleus simultaneously analyzed for single-nucleus ATAC-seq (snATAC-seq) and RNA-seq (snRNA-seq). By introducing strand-tagging into the construction of snATAC-seq libraries, duplex sequencing reduces sequencing error by >10,000-fold while eliminating artifactual mutational signatures. When applied to 98%/2% mixed cell lines, Duplex-Multiome identified sSNVs present in 2% of cells with 92% precision and accurately captured known sSNV mutational spectra, while revealing unexpected subclonal lineages. Duplex-Multiome of > 51,400 nuclei from postmortem brain tissue captured sSNV burdens and spectra across all major brain cell types and subtypes, including those difficult to assay by single-cell whole-genome sequencing (scWGS). This revealed for the first time that diverse neuronal and glial cell types show distinct rates and patterns of age-related mutation, while also directly discovering developmental cell lineage relationships. Duplex-Multiome identified clonal sSNVs occurring at increased rates in glia of certain aged brains, as well as clonal sSNVs that correlated with changes in expression of nearby genes, in both neurotypical and autism spectrum disorder (ASD) individuals, directly demonstrating that somatic mutagenesis can contribute to gene expression phenotypes. Duplex-Multiome can be easily adopted into the 10X Multiome protocol and will bridge somatic mosaicism to a wide range of phenotypic readouts across cell types and tissues.

## Main

Somatic mutations arise in every cell of the human body, but elucidating their role in development, aging, and disease depends on methods that can accurately characterize somatic mosaicism as well as assess their impact on cellular function. While somatic mutations accumulate with age, in patterns and rates that are highly specific to tissues^1–5^, little is known of the extent to which distinct, neighboring cell types in a given tissue age. Currently, identification of patterns of somatic mutagenesis relies on analyzing tissues with uniform cell types and simple clonal patterns^6–8^, or on using scWGS^9,10^ or single DNA molecule sequencing after isolating highly pure cell populations from complex tissues^2,11,12^. To our knowledge, current cell-type-specific profiling methods lack the ability to generate paired epigenomic or gene expression information alongside mutational profiles or remain cost prohibitive, low-throughput or are limited to assaying a small number of loci of interest^13–17^. Computational methods have also been developed to detect somatic mutations from existing single cell (or single-nucleus) RNA-seq (snRNA-seq) and ATAC-seq datasets^18,19^ but remain impossible to apply to complex non-neoplastic tissue due to the low prevalence of true somatic mutations compared to false positives caused by single-stranded DNA damage, PCR and sequencing error. Due to the lack of technology for assessing somatic mutation in a cell-type-specific fashion, the breadth of somatic mutagenesis within complex tissues is likely vastly underappreciated.

### Duplex-Multiome enables low-error sSNV calling from snATAC-seq data paired with snRNA-seq

We selected the 10X Single Cell Multiome ATAC + Gene Expression protocol as a foundation, as this protocol has been widely applied across tissues with success at identifying even rare cell populations^20–24^. However, sequencing errors, with rates ranging from 1 in 10^2^ to 1 in 10^3^ on standard sequencing platforms^25,26^ pose the greatest obstacle to somatic mutation calling in 10X Multiome data, especially in non-malignant tissues in which mutation rates are 1 in 10^7^ or even lower^2,10,11^.

Duplex consensus sequencing, in which products generated from both strands of an initial DNA molecule are sequenced independently, has emerged as the most effective means of reducing sequencing error^2,11,12,25^. We therefore developed Duplex-Multiome, incorporating a strand-tagging strategy to expand the capabilities of the 10X Multiome protocol (Fig. 1a, Extended Data Fig. 1). Duplex-Multiome enables simultaneous detection of somatic single nucleotide variants (sSNVs) and generation of 10X snRNA-seq and snATAC-seq libraries in the same nuclei, addressing the need for high-throughput profiling of cell-type-specific somatic mutational patterns while simultaneously characterizing transcriptomic and epigenomic phenotypes (Fig. 1b).

**Fig. 1.**
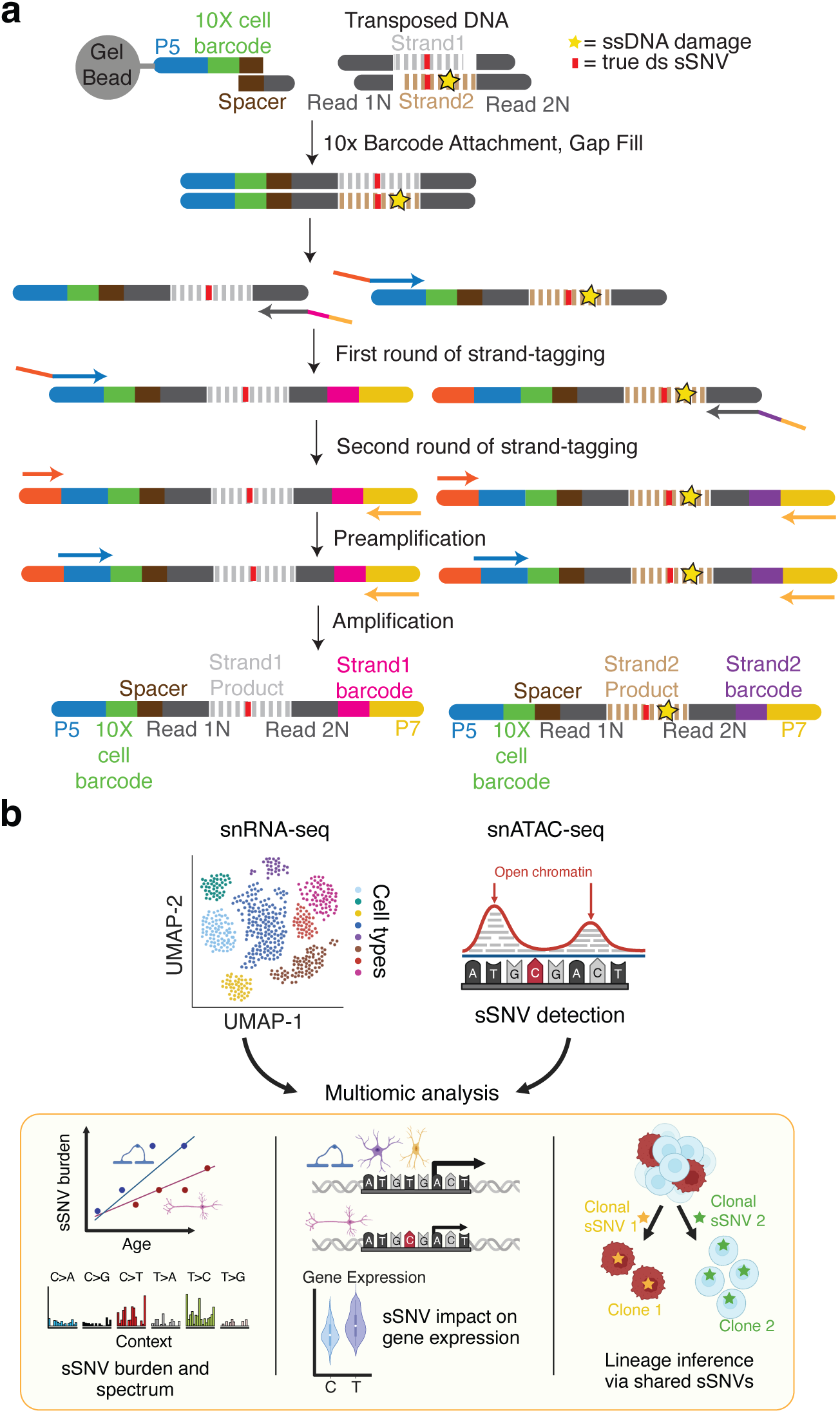
Schematics of Duplex-Multiome protocol and computational pipeline. **a.** Experimental workflow of Duplex-Multiome starting from the GEM (Gel Bead-In Emulsion) Generation & Barcoding step of the 10X Multiome protocol. Steps not shown are identical to the published manual (Methods). Single-stranded DNA damage (yellow star) and true double-stranded DNA single-nucleotide variants (red rectangle) are tracked throughout. **b.** Computational workflow of Duplex-Multiome starting from preprocessed reads generated by Cell Ranger ARC (Methods).

As a pilot, we performed Duplex-Multiome on a human brain sample from a prenatal donor at 21 weeks gestation (Supplemental Table 1). We first examined the lowest error rate, encompassing sequencing, PCR and other sources of error, achievable without requiring duplex consensus, finding that the rate could be reduced down to 10^-5^ by increasing variant calling stringency without regard to strand (Extended Data Fig. 2a, Methods). As this false positive rate still far exceeds known somatic mutation rates in healthy tissues^2,10,11^, we further examined the spectrum of erroneously-called candidate somatic variants and found that they matched a known sequencing error artifact signature, SBS45, from the Catalogue Of Somatic Mutations In Cancer (COSMIC) mutational signatures^27^ (Extended Data Fig. 2b). In contrast, requiring the minimal duplex consensus (i.e., the same variant called on at least one read from each strand) eliminated the artifact signature, reducing error rates down to the extent that could be achieved with even 10-fold sequencing coverage without requiring duplex consensus (i.e., the same variant called on ten or more reads without considering strand). Increasing levels of duplex consensus (e.g., requiring consensus between at least two, three, or more reads from each strand) further reduced error rate (Extended Data Fig. 2a,b).

### Mutation burdens and patterns in mixed cell lines derived from the same donor

We benchmarked Duplex-Multiome using a cell line mixture (1:50 ratio), denoted as COLO829BLT-50, of COLO829 melanoma fibroblast cells (COLO829T or “tumor cells”, Fig. 2a) and COLO829 B-lymphoblast cells (COLO829BL or “BL cells”, Fig. 2a) from the same donor (COLO829), for which known somatic mutations are well-characterized and have been used to assess other somatic mutation detection methods^28^ (Fig. 2a). We performed Duplex-Multiome on COLO829BLT-50 in two replicates, confirming through clustering of multi-omic data along with cell type annotation that two cell types were present, with no detectable batch differences between replicates. Based on cell markers, 4,989 B-lymphoblasts (“BL”, 97.5%) and 128 melanoma fibroblasts (“tumor”, 2.5%) were annotated at the expected ratio of the two cell types in the mixture (Fig. 2b, Extended Data Fig. 3a-c, Methods).

**Fig. 2.**
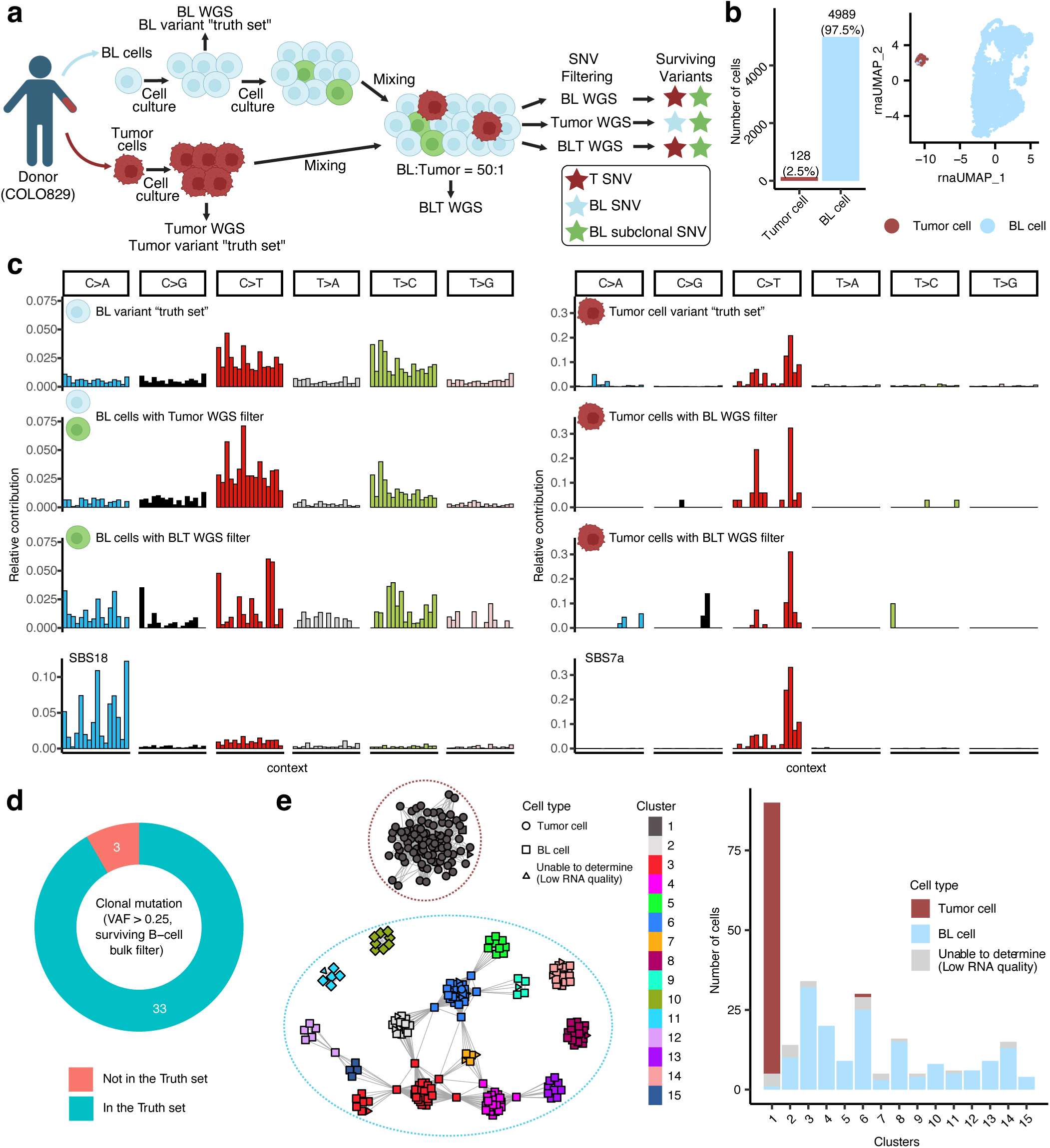
Benchmarking of Duplex-Multiome in COLO829BLT-50 line mixing experiment. **a.** Schematic of the COLO829BLT-50 cell line mixture and strategies for variant filtering. COLO829BL cells (blue BL cells), COLO829T cells (red tumor cells) and subclonal COLO829BL cells (green BL cells) originated from the same donor COLO829. Also see Methods. **b.** Single-nucleus multi-omic profiling of melanoma fibroblast cells (tumor cells) and COLO829BL cells (BL cells). The left panel shows cell type abundances, while the right panel presents uniform manifold approximation and projection (UMAP) coordinates (based on transcriptomic profiles) with annotated clusters (Methods). **c.** Mutational spectra of the COLO829BLT-50 mixture and comparison to COSMIC somatic mutation signatures. The first row shows mutational spectra of truth set variants of BL and tumor cells. The middle two rows display mutational spectra for Duplex-Multiome BL and tumor cells, filtered using bulk WGS from different sources (see panel details). The last row shows COSMIC mutational signatures (SBS18 and SBS7a). **d.** Overlap of Duplex-Multiome clonal sSNVs (VAF > 25%) in tumor cells with the truth set variants that were also called with VAF > 25%. **e.** Inferred lineage relationships between BL and tumor cells. Nodes represent single cells with detected clonal sSNVs, and edges indicate the reciprocal number of shared sSNVs (Methods). Clusters are annotated and colored using label propagation algorithm. Dashed lines circle cell type groups corresponding to the panel **b**, and the bar plot (right panel) shows cell type abundances in each cluster.

“Truth set” SNVs restricted to COLO829T, or COLO829BL, were identified from bulk WGS of each cell line (Methods). In order to determine if Duplex-Multiome could recover these truth set variants, we called sSNVs in the BL and tumor cell populations from our COLO829BLT Duplex-Multiome data and applied a filtering approach consistent with the methods used to generate the truth sets (Methods). As expected, the truth set mutational spectrum for COLO829BL cells was successfully recovered from the COLO829BLT-50 cell line mixture when the COLO829T bulk WGS data was used for germline filtering and vice versa (second row of Fig. 2c, Extended Data Fig. 3d, e).

When bulk WGS from the COLO829BLT-50 cell line mixture was used for germline filtering, the truth set mutational spectrum was also recovered from COLO829T cells (third row of Fig. 2c, Extended Data Fig. 3f). On the other hand, the COLO829BL mutational spectrum was not observed in BL cells under the same COLO829BLT-50 bulk WGS filtering, likely due to COLO829BL cells making up about 98% of the COLO829BLT-50 mixture. Instead, their spectrum resembled a mutational signature for oxidative damage (SBS18) (third row and bottom row of Fig. 2c, left) often seen in cell cultures grown in atmospheric levels of oxygen^29^. These therefore likely represent variants that arose in subclones or individual cells during the cell culture of the COLO829 BL cell line and thus were not captured by bulk WGS. Overall, Duplex-Multiome accurately detected distinct mutational patterns present in clones making up as little as ∼2% of total cells. Our tumor cell Duplex-Multiome clonal variants additionally had high overlap with COLO829T truth set variants (92%; Fig. 2d), further demonstrating the precision of our method.

We investigated whether variants detected by Duplex-Multiome were sufficient to serve as intrinsic barcodes to develop a lineage tree^30^ for COLO829BLT-50 by constructing a graph in which nodes represented cells and the distance between cells represented the reciprocal of the number of sSNVs detected as shared between cells (Fig. 2e, Extended Data Fig. 3g, h). To identify clones and subclones, clusters in the graph were then annotated using an unbiased clustering algorithm (Methods), revealing two main clusters of cells. Cell type annotations derived from our Duplex-Multiome snRNA-seq revealed that most of the tumor cells were located in one large cluster (Fig. 2e, red circle, Cluster 1) separated from the B-lymphoblast-dominant clusters (Fig. 2e, blue circle, Clusters 2-15). Surprisingly, B-lymphoblasts were organized into multiple distinct subclusters (Fig. 2e, Clusters 2-15), defined by variants not present in the COLO829BL truth set, suggesting that sSNVs arising during cell culture defined distinct subclones. Thus, Duplex-Multiome was able to recapitulate the expected clonal structure of COLO829BLT-50 solely based on shared variant information, identify cell-type-specific somatic variants and mutational spectra, as well as disentangle clonal structure of cells making up as little as ∼2% of the total sample from the main lineage.

### Cell-type-specific sSNV accumulation in the aging human brain

The human brain represents the most difficult test for cell-type-specific analysis of somatic mutation, as it is estimated to contain as many as hundreds of distinct cell types^31^, intermingled in a complex, polyclonal architecture^14,16,32–34^. Function-altering, clonally shared somatic mutations in the brain acquired during development have roles in neurodevelopmental disorders^35^, while mutations accumulated after birth have been proposed to play a role in aging as well as neurodegenerative diseases^2,9,36^. We explored the distribution of somatic mutations and their impact across brain cell types across aging by performing Duplex-Multiome on nine neurotypical post-mortem brain samples ranging from 0.2 to 82.7 years of age, generating data for >43,300 nuclei (Fig. 3a, Supplemental Table 1, Supplemental Table 2, Supplementary Fig. 1). Eight of the nine samples also had scWGS data via Primary Template Amplification (PTA)^10^ from neurons available, while five samples had scWGS from both neurons and oligodendrocytes available, all of which served as a “gold standard” for comparison^1^.

**Fig. 3.**
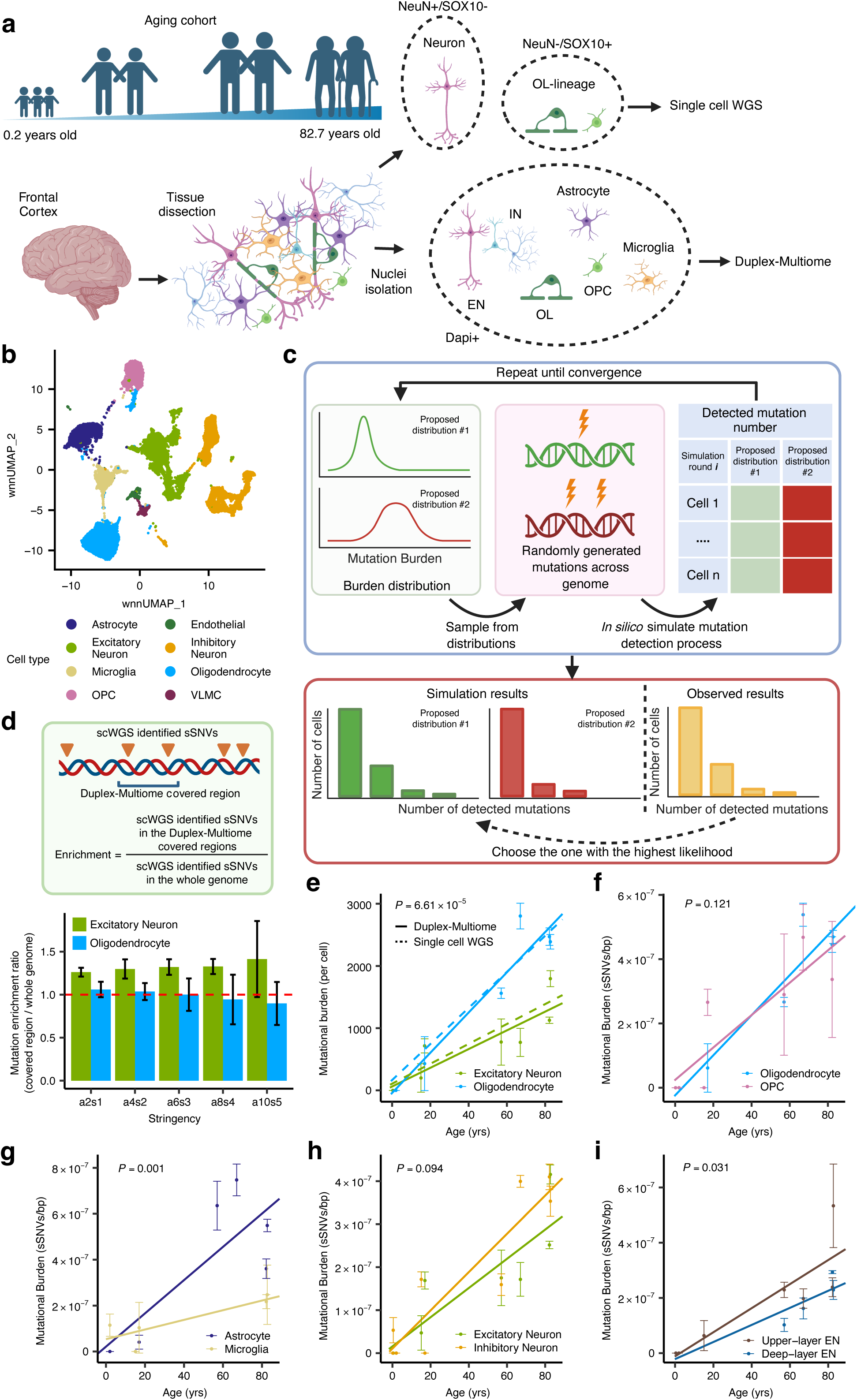
Somatic mutations accumulate in a cell-type-specific manner during brain aging. **a.** Experimental strategy. Nuclei were obtained from the brains of 9 neurotypical individuals through FANS using either DAPI staining (Duplex-Multiome) or staining with NEUN (scWGS neurons) or SOX10 (scWGS oligodendrocytes). Single cell whole genome sequencing (scWGS) was performed using PTA and previously published. SCAN2 was used to call sSNVs from scWGS and our custom pipeline was used to call sSNVs from Duplex-Multiome. **b.** UMAP plot of integrated snRNA-seq and snATAC-seq data using weighted nearest neighbor (WNN) analysis from nine subjects. Cell clusters are annotated and then labelled by colors (Methods). OPC stands for oligodendrocyte precursor cells, and VLMC stands for vascular leptomeningeal cells. **c.** Schematic of the simulation-based maximum likelihood estimation method for mutational burden. Variants within a cell type or a cell cluster are counted to create an observed distribution. A grid search generates candidate mutational burden distributions, and simulations identify the distribution with the greatest likelihood with the observed distribution. This is selected as the mutational burden estimation of the target cluster (Methods). **d.** Distribution of oligodendrocyte (OL) and excitatory neuron (EN) sSNVs in Duplex-Multiome-covered regions versus the whole genome. Enrichment/depletion levels are calculated by comparing sSNV counts in Duplex-Multiome regions to genome-wide counts. These sSNVs are sourced from published scWGS data (Methods). “a2s1”, “a4s2”, “a6s3”, “a8s4”, and “a10s5” refer to regions which are covered by at least 1, 2, 3, 4, and 5 reads from each strand. Error bars indicate ±1 standard deviation across samples. **e – i.** Cell-type-specific sSNV burden in the aging brain. Data points represent the mean sSNV burden per cell type across individuals, with error bars showing the 95% confidence intervals (CI). Trend lines represent fitted linear models, *P* for the difference the difference between cell types in each panel is derived from two-sided two-sample *t*-test (Methods) **e.** Corrected sSNV burden in oligodendrocytes (OLs) and excitatory neurons (ENs), adjusted by Duplex-Multiome-covered region enrichment factors. **f.** Duplex-Multiome sSNV burden in OLs and OPCs. **g.** Duplex-Multiome sSNV burden of astrocytes and microglia. **h.** Duplex-Multiome sSNV burden in ENs and inhibitory neurons (INs). **i.** Duplex-Multiome sSNV burden of EN subtypes: deeper- and upper-layer ENs.

Clustering and cell annotation analysis of the snRNA-seq and snATAC-seq data generated from our Duplex-Multiome experiments confirmed that all expected brain cell types were observed, including astrocytes, excitatory neurons (ENs), inhibitory neurons (INs), microglia, oligodendrocytes (OLs), and oligodendrocyte precursor cells (OPCs) (Fig. 3b, Extended Data Fig. 4a, b).

We examined sSNV burdens across cell types by employing a method that estimates sSNV burden and variance for each cell type by performing a grid search to identify the value pairs with the greatest likelihood based on the observed data (Fig. 3c, Methods). Previous studies have shown differential enrichments of brain sSNVs in open chromatin according to cell type^1,10^. As Duplex-Multiome is expected to be enriched for open chromatin, we verified that neuronal scWGS-identified sSNVs were enriched in regions of neuronal Duplex-Multiome coverage, while oligodendrocyte scWGS sSNVs were slightly depleted in regions of OL Duplex-Multiome coverage (Fig. 3d). To account for this differential sensitivity, we corrected our initial burden estimates by using these enrichment factors for Duplex-Multiome covered regions. Aging curves calculated using this method for excitatory neurons (15.05 ± 3.15 sSNVs/year across the genome, mean ± 1SD) and oligodendrocytes (32.34 ± 2.39 sSNVs/year) were highly concordant with published neuronal and OL scWGS data (Fig. 3e, P = 6.61×10^-5^ for the difference between Duplex-Multiome OLs and ENs, two-sample *t*-test, Methods), supporting that Duplex-Multiome provides cell-type-specific sSNV burdens with high accuracy when open chromatin enrichment factors are known. Furthermore, as Duplex-Multiome sSNV burdens are estimated based on data collected across thousands of cells, the concordance with scWGS data suggests that single-cell mutational burdens—previously defined from small numbers of single cells sequenced genome-wide^2,10^, are indeed characteristic of mutational patterns shared by thousands of cells.

Application of similar methodology to other glial subtypes showed that OLs and OPCs had similar rates of sSNV accumulation ((6.23 ± 0.40) ×10^-9^ and (5.04 ± 1.36) ×10^-9^ sSNVs/bp/year, 35.82 ± 2.30 sSNVs/year and 28.98 ± 7.82 sSNVs/year assuming equal mutation rates across the genome, respectively) (for the difference, *P* = 0.12, two-sample *t*-test, Fig. 3f), suggesting that the increased rate of cell division in OPCs does not greatly contribute to mutational rate, similar to findings in mixed glial cells in the brain and stem cells as well as in differentiated cells in blood and colon^1,2^. However, the reduced number of OPCs in data of older individuals makes it impossible to exclude more subtle increases in sSNV accumulation in OPCs versus OLs. The sSNV accumulation rate differed significantly between astrocytes and microglia, though both accumulate sSNVs at linear rates with age (Fig. 3g, (7.24 ± 1.05) ×10^-9^ sSNVs/bp/year versus (2.10 ± 0.53) ×10^-9^ sSNVs/bp/year, 41.63 ± 6.04 sSNVs/year and 12.08 ± 3.05 sSNVs/year assuming equal mutation rates across the genome, respectively; for the difference, *P* = 0.001, two-sample *t*-test; Fig. 3g). The rate of sSNV accumulation in microglia was surprisingly somewhat lower than that of hematopoietic stem cells with which microglia have a distant shared lineage^37^, but this could reflect low levels of cell division among microglia in brain, or perhaps lower rates of somatic mutagenesis in Duplex-Multiome covered regions, which are enriched for open chromatin.

Turning our attention to major neuronal subtypes, we found that excitatory neurons (ENs) and inhibitory neurons (INs), as annotated by the snRNA-seq component of our Duplex-Multiome both showed linearly increasing sSNV burden with age, with similar rates (3.39 ± 0.76) ×10^-9^ and (4.45 ± 0.73) ×10^-9^ sSNVs/bp/year (15.05 ± 3.15 sSNVs/year and 20.03 ± 3.18 sSNVs/year assuming an equal enrichment rate in covered regions across neuronal cell types) (Fig. 3h, P = 0.094, two-sample *t*-test). In contrast, upper-layer ENs had significantly higher burdens of sSNVs across aging than lower-layer ENs ((4.36 ± 1.01) ×10^-9^ sSNVs/bp/year versus (3.09 ± 0.10) ×10^-9^ sSNVs/bp/year, *P =* 0.03, two-sample *t*-test, 18.44 ± 4.17 sSNVs/year and 13.36 ± 0.43 sSNVs/year assuming an equal enrichment rate in covered regions across neuronal cell types, Extended Data Fig 4c, d), suggesting that upper-layer ENs are more sensitive to neuronal mutagenic processes (Fig. 3i, Fig. 4d).

**Fig. 4.**
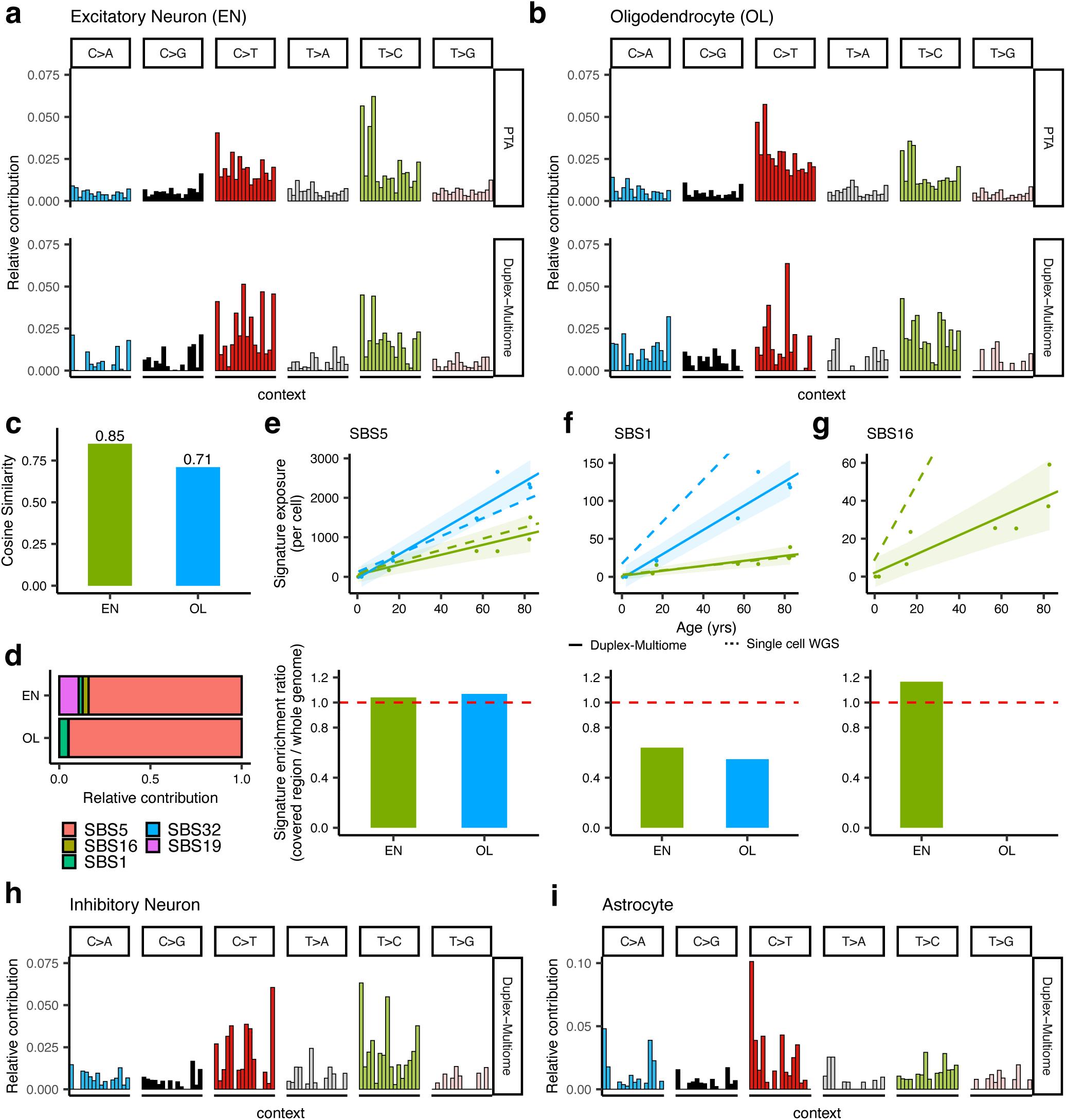
Cell-type-specific mutational spectra and mutagenesis in the aging human brain. **a – b.** Mutational spectra of pooled EN (panel **a**) and OL (panel **b**) sSNVs identified by scWGS with PTA (upper row) and Duplex-Multiome (bottom row). **c.** Cosine similarity of EN and OL between spectra of sSNVs identified by scWGS with PTA and Duplex-Multiome (Methods). **d.** The bar plot shows contributions of the selected COSMIC SBS signature which are known to be present in these cell types (SBS1, SBS5, SBS16, SBS19, and SBS32) to the mutational spectra of EN and OL, calculated by MutationalPatterns (Methods). **e – g.** Distinct SBS signature contributions in open chromatin versus the whole genome. The first row shows the signature exposure levels of SBS5 (panel **e**), SBS1 (panel **f**), and SBS16 (panel **g**) sSNVs identified by scWGS (dotted lines) and Duplex-Multiome (solid lines), which represent signature exposure in open chromatin and the whole genome, respectively. Semi-transparent shading indicates the 95% confidence interval. The second row represents enrichment/depletion levels of SBS signature exposure (Methods). **h – i.** Mutational spectra of pooled IN (panel **h**) and astrocyte (panel **i**) sSNVs identified by Duplex-Multiome.

### Cell-type-specific mutagenesis in the aging human brain

Mutational processes identified in different brain cell types by Duplex-Multiome were largely concordant with those identified from scWGS. EN and OL sSNVs showed a prominent pattern resembling the SBS5 ‘clock-like’ signature (Fig. 4a-c, Extended Data Fig. 5, Supplementary Fig. 2). Fitting our Duplex-Multiome spectra to COSMIC signatures known to be present in these cell types^1^ also revealed contributions of transcription-associated mutational signature SBS16 in ENs and cell division associated signature SBS1 in OLs, as previously observed (Fig. 4d). In contrast to previous studies, SBS19 was detected in ENs but not OLs, although since SBS19 tends to be concentrated in small subsets of cells, this may merely reflect sampling. In addition, while contributions of SBS5 to sSNV burdens recapitulated scWGS results (Fig. 4e), contributions of SBS1 and SBS16 differed somewhat from those previously observed with scWGS (Fig. 4f,g), which may represent differential enrichment of mutagenic mechanisms in open versus closed chromatin.

In order to test whether certain mutational mechanisms may be enriched or depleted in Duplex-Multiome covered regions, we first examined the mutational signature SBS1, a well-characterized mutational signature consisting of C>T variants at CpG sites, which are thought to result from deamination of methylated cytosine^38,39^.

SBS1 has previously been shown to be depleted in open chromatin, likely due to reduced cytosine methylation in active chromatin^1,27^. We examined the distribution of scWGS SBS1 sSNVs and confirmed they are indeed depleted in Duplex-Multiome-covered regions (Fig. 4f, about 2-fold depletion). Consistent with this, Duplex-Multiome detected about a 2-fold reduction in SBS1 in OLs. The magnitude of reduction was consistent with the reduction expected from the distribution of scWGS sSNVs in Duplex-Multiome-covered regions, suggesting that SBS1 variants do not further reduce chromatin accessibility. Therefore, Duplex-Multiome indeed has the capability to detect differential action of mutational mechanisms on open chromatin.

We next turned our attention to whether Duplex-Multiome could illuminate the activity of SBS16, a signature of unknown etiology comprising of mainly A[T>C] variants enriched on the transcribed strand of active genes^27,39^, in open chromatin. Previous studies observed SBS16 predominantly in ENs relative to OLs, with an enrichment in open chromatin^1,10^. We calculated the distribution of scWGS SBS16 sSNVs over our Duplex-Multiome covered regions, confirming that these variants are also expected to be enriched in our Duplex-Multiome data (Fig. 4g). Intriguingly, SBS16 was present in the EN Duplex-Multiome spectrum at a relatively minor contribution, contrasting with the enrichment expected from the distribution of scWGS sSNVs. This observation raises the possibility that SBS16-like A[T>C] sSNVs have the capability of reducing chromatin accessibility, leading to their reduced detection by ATAC-based methods such as Duplex-Multiome.

Duplex Multiome also enabled analysis of mutational spectrums of inhibitory neurons (INs) and astrocytes which have not been previously assayed by scWGS (Fig. 4h,i). The IN spectrum most closely resembled that of ENs while the spectrum of astrocytes most closely resembled that of OLs (Extended Data Fig. 5c-e).

### Analysis of single-stranded DNA damage

Variants present on single strands, representing DNA damage, could be detected in addition to fixed sSNVs detected on both strands. Due to our strand-tagging strategy, ∼1% of strand 2 could acquire two different strand 2 barcodes after the first round of amplification (Extended Data Fig. 6). With the same logic of duplex consensus for double-stranded sSNV calling applied to the same strands, we could distinguish artifacts introduced during later rounds of amplification and sequencing from single-stranded DNA (ssDNA) damage occurring before the first round. This enabled more accurate detection of ssDNA damage (Extended Data Fig. 1, 6, Methods). Across cell types, the spectrum of ssDNA damage events closely resembled COSMIC signature SBS30 (Extended Data Fig. 7), consistent with a recent study which directly sequenced strands of DNA to detect ssDNA damage events^40^. Notably, ssDNA damage showed no correlation with age across cell types (Extended Data Fig. 7a). This, along with the high number of ssDNA damage events detected across samples, suggests they represent DNA damage occurring post-mortem or during library preparation. The strand-tagging strategy of Duplex-Multiome improves discrimination of ssDNA damage events from real sSNVs, again highlighting the utility of our new method.

### Increased rate of clonal variants in glia of aged brains

In addition to “private” sSNVs detected in single cells, which likely represent ongoing mutational processes, Duplex-Multiome also enabled *de novo* detection of clonal sSNVs shared between cells. Clonal sSNVs are of particular interest as their presence across multiple cells makes them potential “retrospective markers” of cell lineage and gives them the greatest likelihood of functional impact. To enhance the detection of such variants, we modified our variant-calling pipeline. Variants present in several cells are likely to be bona fide, since errors occurring at the same genomic position across multiple cells are exceedingly rare, but such real variants may be filtered out due to insufficient supporting reads. Therefore, we optimized the pipeline to include variants shared by three or more cells with at least two supporting reads (denoted as a2s0) as candidate variants. In order to avoid excluding somatic variants arising during early developmental stages, we also relaxed our germline variant filtering criteria. In order to avoid misidentifying germline mutations as somatic, we introduced a Bayesian filtering approach using the Markov Chain Monte Carlo method as an additional layer of germline variant filtering (Fig. 5a). In addition, we applied another filter to avoid systematic errors from sequencing and alignment (Methods). We verified our clonal mutational calling pipeline by testing it on the BLT-50 cell line mixture (Extended Data Fig. 8).

**Fig. 5.**
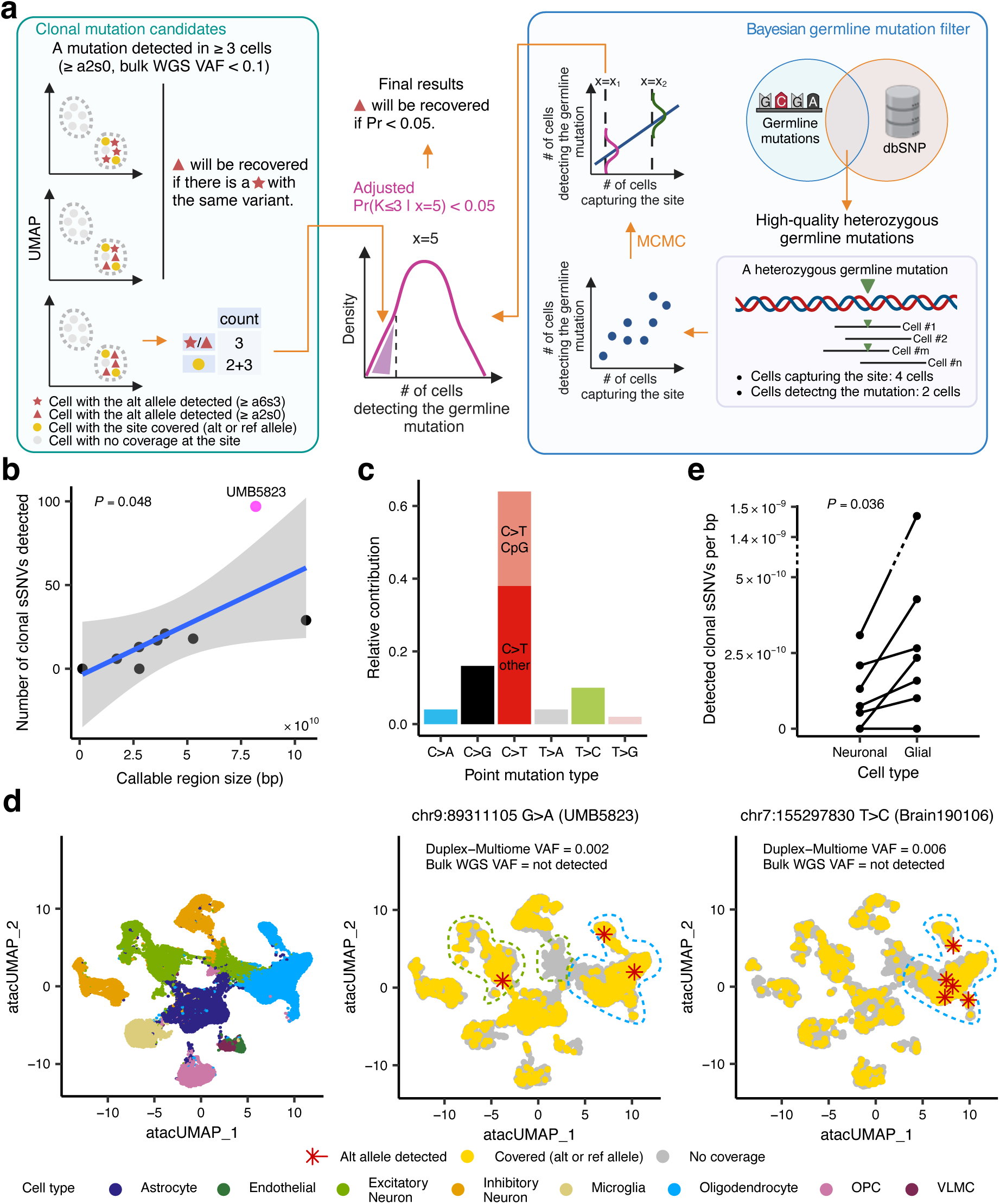
Characterization of clonal sSNVs in the aging human brain. **a.** Computational pipeline for clonal sSNV identification. Variants shared by three or more cells with at least two supporting reads (a2s0) are selected as candidates, because errors occurring at the same position across multiple cells are rare. Our germline variant filtering criteria are relaxed to include early developmental variants. An additional germline filter is applied to exclude genuine germline mutations (Methods). **b.** Relationship between the number of clonal sSNVs detected in each sample and the callable region size. Each dot represents an individual, with a linear model trend line fitted to the data. *P* is derived from the linear model. **c.** Base substitutions of clonal sSNVs across all subjects. **d.** UMAP plot of snATAC-saq data showing examples of clonal sSNV. Left: UMAP plot of cells including recovery of cells with low-quality transcriptome data but good-quality ATAC data (Methods). Middle: G>A variant at chr9:89311105 in sample UMB5823, found in two OLs and one EN. The variant has a VAF of 0.0016 in Duplex-Multiome and is not detected in 30X bulk WGS due to low VAF. This variant was validated by Amplicon-seq (Methods) with VAF of 0.0018. Right: T>C variant at chr7:155297830 in sample Brain1901106, found only in the OL lineage. Detected in five OLs with a VAF of 0.006 in Duplex-Multiome but absent in 30X bulk WGS. Red stars indicate cells where the variant (alt allele) was detected by Duplex-Multiome. Yellow dots represent cells where the variant site was covered by Duplex-Multiome (either alt or ref allele could be detected). Grey dots denote cells where the variant site was not covered by Duplex-Multiome. **e.** Difference of the clonal sSNV rates between neurons (EN and IN) and glial cells (astrocyte, oligodendrocyte, microglia, and OPC). Each sample is shown by a pair of dots, one for glial cells and one for neurons, connected by a line. Samples by the order of glial cell clonal sSNV rate from highest to lowest are: UMB5823, Brain190106, UMB1864, UMB5657, UMB1278, UMB5451, UMB1465, UMB4428, and UMB4638. No clonal sSNVs were called in UMB1465, UMB4428, or UMB4638 after filtering. Their points therefore overlap at 0. *P* is calculated with a paired two-sided Wilcoxon test (3.4-fold higher in glial cells).

Duplex-Multiome detected up to 27 clonal sSNVs per sample, with the count being significantly proportional to the callable region size (Fig. 5b, P = 0.048, linear regression). We validated 45 of these clonal sSNVs via Amplicon-Seq, including variants in all samples for which we still had tissue available. The validation rate of our clonal sSNVs was above 90% for sSNVs with variant allele fraction (VAF) of 0.01 or greater as calculated by either method (9 out of 10 for Duplex-Multiome VAF ≥ 0.01, 6 out of 6 for Amplicon-Seq VAF ≥ 0.01; Supplemental Table 3). This confirms that Duplex-Multiome has high precision in detecting clonal sSNVs that occurred early in development. Interestingly, most clonal sSNVs detected by our pipeline had VAFs of below 0.01 (38 out of 50 with Duplex-Multiome VAF < 0.01, 39 out of 45 with Amplicon-Seq VAF < 0.01; Extended Data Fig. 9a-c; Methods). As the current “gold-standard” for *de novo* clonal sSNV discovery in the brain, ultra-deep WGS, has a detection threshold of ∼0.02 while Duplex-Multiome detects clonal variants with VAFs of down to 0.001 (Supplemental Table 3), this suggests that Duplex-Multiome can discover clonal variants that are an order of magnitude rarer than current methods^36,41^. Indeed, Amplicon-Seq validated 71% of these rarer variants (25 out of 35 for Duplex-Multiome VAF < 0.01, 28 out of 39 for Amplicon-Seq VAF < 0.01). A lower validation rate for sSNVs with VAFs of less than 0.01 is expected, as such variants may be focally restricted and therefore difficult to recapture in subsequent samplings, even from the same tissue^14,30^. Along with cell-type-specific chromatin accessibility, this sampling issue could also explain discordance in VAFs in Duplex-Multiome versus Amplicon-Seq for these rarer variants. Clonal sSNVs detected by Duplex-Multiome had a greater proportion of C>T variants at CpGs (Fig. 5c), recapitulating previous findings of clonal variation in the human brain occurring due to mutagenesis in dividing progenitors^1,12,33,36^.

Duplex-Multiome’s unique ability to detect clonal sSNVs while simultaneously identifying cell types enabled direct identification of cells sharing sSNVs (Fig. 5d) as well as direct comparison of clonal sSNV rates in neuronal versus glial cells. When clonal sSNVs of all VAFs were included, we did not observe a significant difference in clonal sSNV rates between neurons and glia (*P* = 0.18), likely because these include high-VAF sSNVs which occurred early in development and are therefore dispersed across neuronal and glial lineages^33,36^. We therefore focused on low-VAF clonal sSNVs (VAF < 0.01) and discovered that glial cells had a significantly higher rate of such clonal sSNVs than neurons (*P* = 0.036; Fig. 5e). In addition, an 82.7-year-old sample UMB5823 had a higher clonal sSNV rate than our other samples, including another 82-year-old (Fig. 5b), particularly in glial cells (*P* = 0.027, linear regression; Extended Data Fig. 9d). These results are consistent with the accumulation of somatic mutations in aging glial progenitors in certain samples, which are inherited by daughter cells across aging, in contrast to post-mitotic neurons which can only share somatic mutations accumulated during development. Previous studies examining clonality in human brains had difficulty determining whether higher numbers of clonal sSNVs in aged brain samples were due to increased clonal sSNVs in glial cells or due to infiltrating blood cells^41^. Our results support that such clonal sSNVs indeed arise in aging glia without substantial contribution from blood cell clones, at least in healthy brain samples. We also observed a 67-year-old sample, Brain190106, had a similar glial-to-neuronal ratio of clonal sSNVs as UMB5823 (Fig. 5e). Brain190106 also carried one sSNV, which was only detected in the OL-lineage (Fig. 5d; chr7:155297830 T>C, 5 OLs with the variant, 823 covered cells). Together, these results suggest that clonal sSNVs frequently arise in glial cells, whose lineage can be disentangled by Duplex-Multiome.

### Developmental, non-coding somatic mutations correlating with changes in gene expression in neurotypical and ASD brains

Since Duplex-Multiome simultaneously detects sSNVs and performs snRNA-seq, our method is ideally poised to assay directly the functional effects of somatic mutation on gene expression. We developed an empirical statistical model using bootstrapping to compare gene expression in cells carrying a sSNV versus those cells in which the sSNV was not detected, taking allelic dropout into account. We limited our pipeline to assessing differential gene expression in a 1Mb interval around each variant, positing that variants in regulatory regions would most likely affect gene expression in *cis*, affecting genes nearby on the same chromosome (Fig. 6a, Methods). In order to test our approach, we performed Duplex-Multiome on an ASD sample AN06365 (Supplementary Fig. 3a) previously found to carry a clonal somatic T>C variant in a brain enhancer region (chr1:205284910 T>C) that reduces enhancer activity^36^. We detected cells of all expected types by the snRNA-seq component of Duplex-Multiome (Fig. 6b-d). Presence of chr1:205284910 T>C correlated with significantly reduced expression of the nearby *NUCKS1* gene (*P* = 0.025) within AN06365 cells. We also observed a small but statistically significant decrease in *NUCKS1* expression in AN06365 compared to 9 neurotypical samples without the T>C variant (Fig. 6e, P = 3.1×10^-4^). This reduction may result from the presence of mutant cells. These findings support that somatic mutation can indeed result in changes in gene expression and that Duplex-Multiome is capable of detecting such changes.

**Fig. 6.**
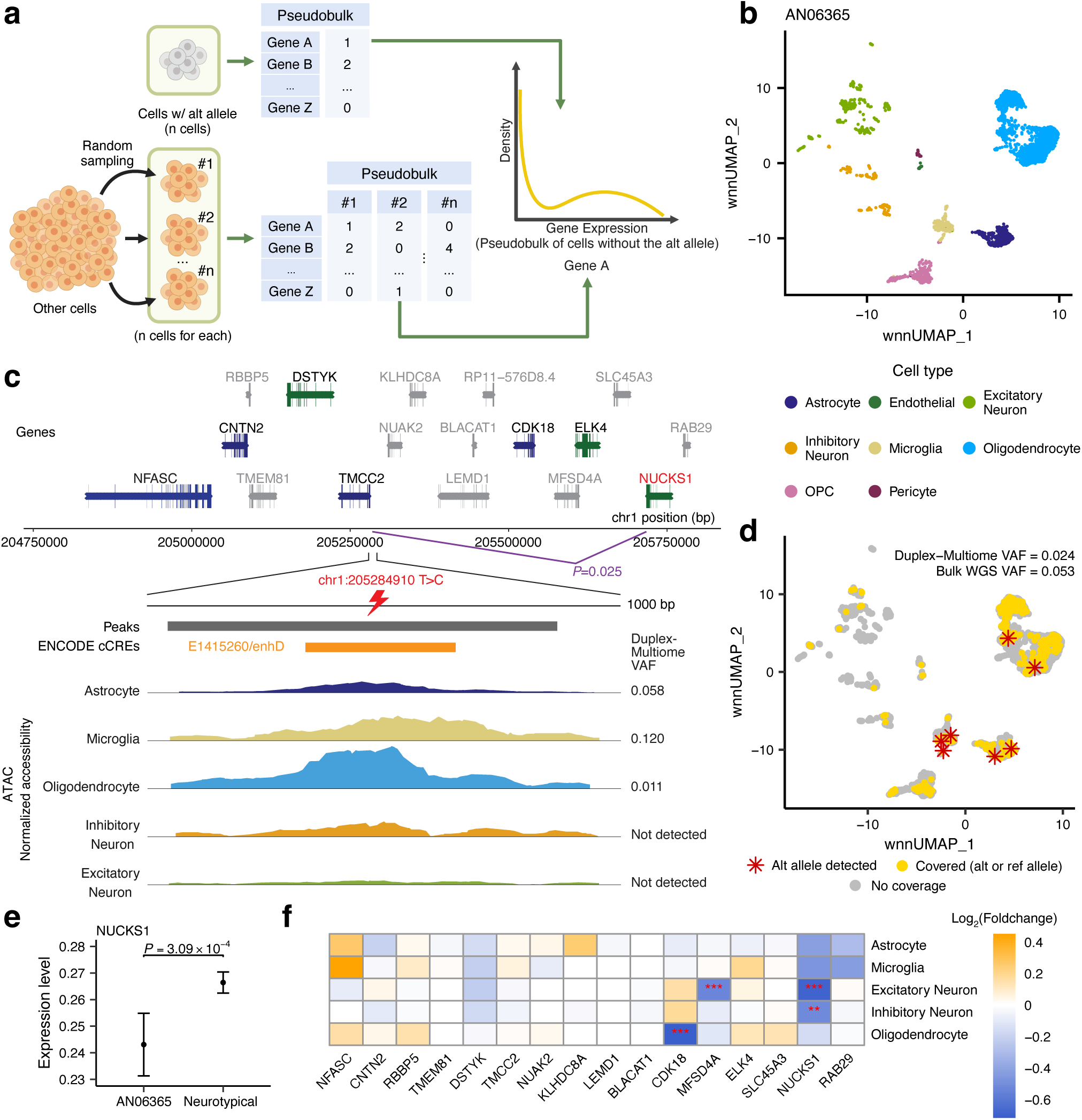
An ASD brain harbors non-coding clonal sSNVs associated with changes in gene expression *in cis*. **a.** Computational pipeline for discovering association between non-coding clonal sSNVs and nearby gene expression. Using an empirical statistical model with bootstrapping, gene expression in cells carrying a sSNV is compared to cells in which the sSNV was not detected. The analysis focuses on genes within a 1Mb interval around each variant (Methods). **b.** UMAP plot of integrated snRNA-seq and snATAC-seq data using WNN analysis from the ASD brain sample (AN06365). OPC stands for oligodendrocyte precursor cells. **c.** Analysis of the T>C variant at chr1:205284910 in AN06365. Genes within a 1Mb interval of the variant are shown, with those expressed in fewer than 10% of cells excluded (colored grey). Blue and green indicate direction of transcription of analyzed genes. Peaks from snATAC-seq data and ENCODE cCREs are displayed. The bottom panel shows normalized snATAC-seq signals across the region and Duplex-Multiome VAF of this variant in five main brain cell types. *P* is calculated with a one-sided empirical test (Methods). **d.** UMAP plot showing the T>C variant from AN06365. Duplex-Multiome detected the variant with a VAF of 0.024 and the previous bulk WGS detected it with a VAF of 0.053. Red stars indicate cells where the variant (alt allele) was detected by Duplex-Multiome. Yellow dots represent cells where the variant site was covered by Duplex-Multiome (either alt or ref allele could be detected). Grey dots denote cells where the variant site was not covered by Duplex-Multiome. **e.** *NUCKS1* expression in AN06365 compared to neurotypical brains. Data points represent mean expression values, with error bars indicating the 95% confidence interval across all single cells in indicated sample(s). *P* is calculated with a one-sided Wilcoxon test for expression in AN06365 and the neurotypical brains across single cells (about 8% decrease in *NUCKS1* expression in AN06365). **f.** Heatmap showing log2(fold changes) of nearby genes between AN06365 and another ASD sample (ABNCR6D) that does not carry this variant. We calculated *P* for genes with Wilcoxon tests and performed multiple testing correction using Bonferroni correction. *** represents adjusted *P* < 0.001 and ** represents adjusted *P* < 0.01.

Notably, chr1:205284910 T>C was detected by Duplex-Multiome in OLs, astrocytes, and microglia, suggesting it occurred extremely early in development (likely prior to neurulation, given the early divergence of the microglial lineage) yet was surprisingly not detectable in neurons (Fig. 6c, d). However, our Amplicon-Seq confirmed that chr1:205284910 T>C was present in about 6% of NeuN-sorted neurons as well as in non-neuronal cells (Supplemental Table 3), consistent with it occurring early in development, but suggesting that the lack of detection in neurons could result from a cell-type-specific loss of chromatin accessibility. Supporting this, ATAC-normalized accessibility around the variant site is lowest in neurons, particularly ENs (Fig. 6c). Further examination revealed reduced neuronal chromatin accessibility even in brain samples that did not carry the variant (Supplementary Fig. 3b, c), suggesting that this genomic region has inherently reduced chromatin accessibility in neurons. We wondered whether this meant that this previously-characterized brain enhancer region was primarily active in non-neurons and therefore if the variant would primarily impact gene regulation in non-neuronal cells.

In order to enable gene expression comparison between cell types and control for ASD as a covariate, we turned to differential gene expression analysis between AN06365 and another ASD sample, ABNCR6D, which did not carry the variant. Our snRNA-seq data showed reduced *NUCKS1* expression across all major brain cell types, with the reduction being statistically significant in ENs and INs (Fig. 6f). Interestingly, *CDK18* was downregulated specifically in OLs while *MFSD4A* was downregulated specifically in ENs, suggesting that the same clonal sSNV might impact gene expression in cell-type-specific ways, and that Duplex Multiome can detect this.

However, as differential expression of these genes was observed only when considering all cells of each type, regardless of known variant status, we cannot exclude whether other properties of AN06365 contribute to these gene expression changes. Nevertheless, these results indicate that gene expression across both neurons and non-neurons is impacted in AN06365, despite low chromatin accessibility in neurons. Duplex-Multiome thus sheds light on the potential impact of chr1:205284910 T>C across cell types, highlighting the utility of Duplex-Multiome’s paired snRNA-seq and cell type information.

We next expanded our functional impact analysis to determine if we could detect clonal sSNVs correlating with changes in gene expression in neurotypical brain samples. We detected one G>A variant (chr2:19990244 G>A) located in a promoter in a 15.1-year-old brain sample, which correlated with upregulation of the gene *LAPTM4A* ∼60kb away via both comparison of cells detected with the variant versus cells without within the sample, as well as gene expression analysis across samples, as described above (*P* = 0.006, Extended Data Fig. 10a-d). We cloned constructs containing mutant and reference promoter sequences and performed a luciferase assay in a neuroblastoma cell line, confirming that the sequence has increased activity over a minimal promoter, with a trend towards further increased activity by the mutant promoter over the reference (Extended Data Fig 10e, *P* = 0.057). This suggests that the variant may lead to upregulation of nearby genes via increasing promoter activity. Together, these observations suggest that non-coding somatic mutations in both neurotypical as well as ASD brains can contribute to changes in gene expression. Clonal sSNVs thus represent an intriguing source of gene expression variation between individuals to investigate in future studies.

## Discussion

Integrating strand-tagging with 10X Single Cell Multiome ATAC + Gene Expression protocol, we expanded the capabilities of this widely-used 10X protocol to enable highly accurate somatic variant calling alongside gene expression and epigenetics in a high-throughput manner. Our new Duplex-Multiome method is particularly relevant for the study of somatic mutations in complex tissues, such as the human brain, in which cell types of interest are highly intermingled and laborious to isolate.

Previous studies of somatic mutagenesis in the human brain have been mainly restricted to neurons, though glial and neuronal subtypes are increasingly appreciated to play roles in both typical development and aging, as well as neurodevelopmental disorders, brain cancer, neurodegeneration. In addition, the study of functional impact of somatic mutations has primarily focused on coding mutations that directly alter amino acid sequences, despite the majority of somatic mutations occurring in non-coding regions. Duplex-Multiome bridges these gaps in knowledge by characterizing sSNVs in all major brain cell types and sub-types across aging, as well as by elucidating how different types of sSNVs impact chromatin accessibility and gene expression.

Duplex-Multiome reveals, for the first time to our knowledge, that upper layer ENs and astrocytes are the EN and glial cell types most susceptible to somatic mutagenesis, respectively. While the limited coverage of the method renders genome-wide mutational patterns difficult to definitively determine, the focused sequencing on open chromatin also enables the first direct assessment of the functional impact of somatic mutagenesis on epigenetic state in the human brain. In particular, we identify the cell-division-associated signature SBS1, and the transcription-associated signature SBS16 as depleted in open chromatin, with the latter appearing to reduce chromatin accessibility in itself. We also discovered cases of somatic mutations correlating with changes in nearby gene expression, as well as frequent clonal sSNVs in glial cells.

Previous studies showed that elevated somatic mutation burden does not result in premature aging in healthy cells^42^. In agreement with this, most variants we detected did not have functional impact. Our findings suggest that specific somatic mutations, rather than overall mutation burden, may drive alterations in chromatin state and gene expression.

While this study focused on the brain, Duplex-Multiome can easily be applied to any system in which the 10X Multiome has been successfully utilized, enabling somatic mutation detection to be added to any multi-omic analysis. Other complex tissues consisting of highly intermingled cell types, such as the kidney, would be particularly relevant. Widespread application of Duplex-Multiome to hundreds and thousands of single nuclei will provide clearer insights into findings currently limited by sparsity. In addition, our strand-tagging strategy can in theory be integrated into any single-cell library construction method utilizing tagmentation. Thus, this study paves the way to bridging somatic mutation detection with a myriad of single-cell methods, including those that assay chromatin marks^43^, 3D genome structure^44^, as well as spatial single-nucleus methods^45,46^.

## Methods

### Human tissue samples

Post-mortem human tissues were obtained from the NIH Neurobiobank at the University of Maryland Brain and Tissue Bank, the Boston University UNITE or VA-BU-CLF Brain Bank and the Autism Brain Net according to their institutional protocols. Research on these de-identified specimens and data was performed for this study with approval from the Boston Children’s Hospital Institutional Review Board.

### Bulk whole-genome sequencing read-mapping and generation of BAM files

Matched reference whole genome sequencing (WGS) was required for somatic variant detection by our Duplex-Multiome pipeline. Bulk genomic DNA was extracted using QIAGEN QIAamp DNA Mini, QIAGEN EZ1 kit, or phenol-chloroform extracted and sequenced by either Illumina HiSeq 2000, HiSeq 2500, HiSeq X, NovaSeq 6000, or NovaSeqX to a target mean coverage of 30-45X for all samples except UMB1465 and ABN_CR6D which were sequenced to a target mean coverage of 200X. Bulk WGS data for all brain samples except UMB1465 and ABN_CR6D were previously published^1,47^ and aligned to the reference genome GRCh37 with decoy chromosomes. To realign them to GRCh38, these published BAM files were first converted to FASTQ files containing only well-paired reads. We then performed alignment using BWA v0.7.8^48^ with the reference genome GRCh38. Data preprocessing followed the GATK Best Practices (https://gatk.broadinstitute.org/hc/en-us/articles/360035535912-Data-pre-processing-for-variant-discovery).

Intermediate BAM files from the alignment were sorted using the SortSam function of Picard v2.26.10, and duplicated reads were marked using MarkDuplicates from Picard v2.26.10. Base quality score recalibration was then performed using Genome Analysis Toolkit (GATK) v4.3.0.0.

For the COLO829 cell line, BAM files of bulk whole-genome sequencing data (aligned to GRCh38) were downloaded from the Somatic Mosaicism Across Human Tissues (SMaHT) data portal (https://data.smaht.org/data/benchmarking/COLO829).

### Nuclei isolation and sorting

Nuclei were isolated from the post-mortem human brain samples as previously described with minor modification^49^ (Nuclei Isolation from Complex Tissues for Single Cell Multiome ATAC + Gene Expression Sequencing). Briefly, about 50mg of tissue was scraped from a larger piece of tissue and resuspended in 1mL homogenization buffer with additives (10mM Tris Buffer pH 8.0, 250mM Sucrose, 25mM KCl, 5mM MgCl2, 0.1% Triton X-100, 0.1mM DTT, 1X cOmplete™, Mini, EDTA-free Protease Inhibitor Cocktail (Roche 11836170001), 25ul Protector RNAse inhibitor (Roche 3335399001, 0.2U/µl)) and transferred to a 7mL douncer and dounced 10 times with a ‘tight’ pestle. Homogenized nuclei were spun for 10min at 900g at 4C, then washed once with Blocking Buffer (1X PBS pH 7.4, 1% BSA), spinning for 5min at 400g at 4C. All spins were done in a bucket centrifuge. Nuclei were resuspended in 500ul Blocking Buffer with Protector RNAse inhibitor (Roche 3335399001, 1U/ul) and Dapi (final concentration 1ug/ml) and passed through a 40µm filter. Dapi positive nuclei (50,000 nuclei per 10X reaction) were Fluorescence-Activated Nuclei Sorted (FANS) into 300ul of Lysis Dilution Buffer (10mM Tris-HCl pH 7.4, 10mM NaCl, 3mM MgCl2, 1% BSA, 1mM DTT, 1U/ul RNAse inhibitor) in LoBind Eppendorf tubes prewashed with Lysis Dilution Buffer.

For neuronal sorting, nuclei were isolated following the procedure above up to the douncing step, and then stained in Blocking Buffer with 1:1000 anti-NeuN antibody directly conjugated to Alexa Fluor 488 (Millipore, MAB377X, clone A60, AlexaFluor-488 conjugated) for 20 minutes before resuspension with Blocking Buffer with Dapi. NeuN staining produced a bimodal signal distribution, distinguishing NeuN+ and NeuN-nuclei. 141,000 NeuN+ and NeuN-nuclei were sorted into 60ul of 6X tissue lysis buffer (60mM Tris-HCl, 30mM EDTA, 300mM NaCl, 3% SDS) and genomic DNA purified via phenol-chloroform purification for amplicon sequencing (see below).

### Duplex-Multiome library preparation and sequencing

In order to permeabilize sorted nuclei, 50ul of 1X Lysis Buffer was added (10mM Tris-HCl pH 7.4, 10mM NaCl, 3 mM MgCl2, 0.1% Tween-20, 0.1% Nonidet P40 Substitute, 0.01% digitonin, 1% BSA, 1mM BSA, 1U/ul RNAse inhibitor) to sorted nuclei in 300ul of Lysis Dilution Buffer (10mM Tris-HCl pH 7.4, 10mM NaCl, 3mM MgCl2, 1% BSA, 1mM DTT, 1U/ul RNAse inhibitor). Nuclei were incubated for 2min on ice and then 5ul of 10% Tween-20 was immediately added. Nuclei were then immediately centrifuged at 500g for 5min at 4C. After spinning, all but 5ul of supernatant were removed and 200ul of 1X Diluted Nuclei Buffer (10X Multiome kit) was added. Nuclei in 1X DNB were centrifuged at 500g for 5min and all but 7-10ul supernatant was removed. Nuclei were resuspended in 1X DNB by gentle tapping and 2ul were taken for nuclei counting in a hemacytometer. Concentrations were adjusted to 2000 nuclei/ul in 1X DNB and 5ul of nuclei loaded according to Step 1.0 (Transposition) of the Chromium Next GEM Single Cell Multiome ATAC + Gene Expression manual. Transposition, GEM Generation and Barcoding, and Post GEM Incubation Cleanup were performed according to the manual until Step 3.2j, where 33.4ul Qiagen EB was used to elute purified DNA from SPRIselect beads.

Strand-tagging was performed in place of Step 4 (Pre-Amplification) of the manual as follows: 33ul of the eluted sample, 37ul of pre-amplification enzyme mix (10X PN 2000270/2000274), 1ul of 100uM PBS_P5 (AACTAGGCGTAAGCTGGAGATCGTAAATGATACGGCGACCACCGAGATCTACAC), 1ul of one well of Sample Index N, Set A (10X PN 3000427), and 2ul of 100uM Partial TSO (AAGCAGTGGTATCAACGCAGAG) were mixed by pipetted and spun down briefly. The first round of strand-tagging was then performed in a thermocycler as follows: 72C for 5min, 98C for 3min, 63C for 30s, 72C for 1min, 4C hold. Samples were then treated with 2ul thermolabile ExoI (NEB M0568S) to degrade the remaining primers and incubated in a thermocycler at 37C for 15min, 65C for 15min, 4C hold. The second round of strand-tagging mix was then prepared by adding 1ul of 100uM PBS_P5, 1ul of another well Sample Index N, Set A (10X PN 3000427) [NOTE: Must be a different well than the well chosen for the first round of strand-tagging], 2ul of 100uM Partial TSO, and ul of pre-amplification enzyme mix (10X PN 2000270/2000274). The second round of strand-tagging was then performed in a thermocycler as follows: 98C for 3min, 63C for 30s, 72C for 1min, 4C hold. Samples were then treated with 2ul thermolabile ExoI (NEB M0568S) to degrade the remaining primers and incubated in a thermocycler at 37C for 15min, 65C for 15min, 4C hold. Pre-amplification mix was prepared by adding 1ul of 100uM PBS (AACTAGGCGTAAGCTGGAGATCGTA), 1ul of 100uM P7 (CAAGCAGAAGACGGCATACGAGAT), 1ul of 100uM Partial TSO, and 1ul of 100uM 10X Partial Read 1 R (ACACTCTTTCCCTACACGACGCTC), and 7ul of pre-amplification enzyme mix (10X PN 2000270/2000274). Pre-amplification was then performed in a thermocycler as follows: 98C 3min, 98C 20s, 63C 30s, 72C 1 min, loop back to step 2 6 times, 72C 1min, 4C hold. Pre-Amplification SPRI Cleanup was then performed according to the manual (Step 4.3). In place of Step 5.0 (ATAC Library Construction) and 5.1 (Sample Index PCR), the final amplification mix was prepared by adding 40ul of pre-amplified sample, 50ul of Amp Mix (10X PN 2000047/2000103), 7.5ul SI-PCR Primer B (10X PN 2000128), and 2.5ul of 100uM P7 only primer. PCR was then performed in a thermocycler according to Step 5.1d of the manual. All steps for GEX Library construction were performed as described in the manual.

Duplex-Multiome single-nuclei ATAC-seq libraries were sequenced by Novogene on 1-2 lanes of NovaSeqX (10B flowcell) each, resulting in 1.3 billion to 2.6 billion PE150 reads. Duplex-Multiome single-nuclei RNA-seq libraries were sequenced by Novogene on the NovaSeqX (10B flowcell) according to the recommendations of 10X Genomics.

### 10x single nucleus multiome data analysis

Duplex-Multiome reads were processed using cellranger-arc count v2.0.1^50,51^ for alignment, filtering, barcode counting, peak calling, and quantification of snATAC-seq and snRNA-seq molecules, using the GRCh38 reference genome and the default cellranger parameters. The Multiome data were then analyzed with Seurat v4.0.5^52^ and Signac v1.5.0^53^ under R v4.0.2^54^ (all subsequent analyses in this work were conducted with R v4.0.2 if using R environment). We used the following QC metrics: nCount_ATAC > 1000, nFeature_RNA > 200, and percent.mt < 10%. Cells passing these filters were subjected to doublet removal using scds v1.4.0^55^, where the upper 5% of doublet scores were filtered out.

Next, snATAC-seq and snRNA-seq data from each sample within the same group were combined (two replications of COLO829BLT50, nine healthy brain samples, and two ASD samples). The merged snRNA-seq data were processed by SCTransform, followed by RunPCA, while the merged snATAC-seq data were analyzed with RunTFIDF, FindTopFeatures, and RunSVD. Harmony was used separately for the snRNA-seq and snATAC-seq data to integrate data from each sample and correct batch effects. The integrated data were then analyzed using FindMultiModalNeighbors, RunUMAP, and FindClusters for dimensionality reduction, visualization, and clustering.

For COLO829BLT-50, cell types were annotated using established markers for BL cells and melanoma tumor cells. For all brain samples, cell type labels were first transferred from curated brain scRNA-seq data from the Allen Brain Map^56^ (https://portal.brain-map.org/atlases-and-data/rnaseq/human-multiple-cortical-areas-smart-seq) and manually verified based on markers (https://celltypes.brain-map.org/rnaseq/human_ctx_smart-seq?selectedVisualization=Heatmap&colorByFeature=Cell+Type).

Additionally, we performed sub-cell-type annotation for neurons from the healthy brain samples. We selected clusters annotated as excitatory and inhibitory neurons and generated two new Seurat objects for excitatory and inhibitory neurons, respectively. Data normalization, integration, dimensionality reduction, and clustering were repeated with the same tools. Subtype annotation was based on established neuron subtype markers, and suspicious clusters with expression of markers from multiple subtypes were removed.

### Generating the truth set sSNVs for COLO829BLT-50

For the melanoma tumor cells (COLO829T) from COLO829BLT-50, its truth set was downloaded from the SMaHT data portal (https://data.smaht.org/data/benchmarking/COLO829#truthset). For BL cells (COLO829BL), we used GATK’s Mutect2 to analyze bulk WGS data from BL cells, using the bulk WGS data from tumor cells as a reference. This approach filtered out most germline variants from the patient (COLO829), variants that arose during early developmental stages of the patient, and any potential contamination from the tumor cells. Additionally, high-confidence germline variants for BL cells obtained from SMaHT and a panel of normals from GATK (https://console.cloud.google.com/storage/browser/_details/gatk-best-practices/somatic-hg38/1000g_pon.hg38.vcf.gz) were used to further exclude germline variants and artifacts. This process yielded SNV candidates for BL cells, which occurred during the patient’s aging and following cell culture. These candidates were filtered with a VAF > 0.25 and sequencing depth (DP) > 100 to ensure only high-confidence SNVs were included in the truth set.

### Calling of sSNV candidates from Duplex-Multiome data

The BAM file of processed ATAC reads from Cell Ranger^50,51^ (atac_possorted_bam.bam) was first split by cell and strand barcode using *samtools* v1.21^57^. After adding the corresponding strand tags to the BAM files, the two BAM files (strand 1 and strand 2) for each single cell were combined. Read families (i.e., all reads originating from the same ATAC-tagmented fragment) were annotated based on their start and end positions using the *annotateReadFamilies* function from DuplexTools v17201a76d0c3.

To reduce the risk of misalignment and bacterial DNA contamination, the following reads were excluded for calling: reads with indels, reads located in predefined blacklist regions from published studies, 10 bases at both ends of all reads, and read families shorter than 100 bases. Using *mcsCallVariants* from DuplexTools, germline and somatic mutations were identified with the following criteria: base quality > 30, MapQ > 20, supporting reads ≥ 6 (with ≥ 3 from strand 1 and ≥ 3 from strand 2, denoted as a6s3), and a read family variant allele frequency (VAF) of 1.

Matched bulk WGS data were used to filter out germline mutations, saving only those calls that were absent in any reads from the matched bulk WGS data as sSNV candidates. Additionally, contiguous mutation calls within a read family, which could indicate artifacts, were removed.

Contamination of duplex sequencing libraries with DNA from other individuals could introduce artifacts and overestimate mutational burdens, mainly because germline mutations of other samples will appear as somatic mutations. Therefore, we filtered out variants that were present in the bulk WGS data of other samples in this study, as well as those found in GnomAD v3.1.2 with a variant allele frequency (VAF) > 1% in the population. Additionally, shared somatic mutation calls across samples were discarded.

### Duplex-Multiome covered region sSNV enrichment analysis

Published single-cell WGS data of excitatory neurons and oligodendrocytes were downloaded (https://zenodo.org/records/7508803) for downstream analysis^58^. As described earlier, read families that passed all filters and contained more than 6 reads (with ≥ 3 from strand 1 and ≥ 3 from strand 2, denoted as a6s3) were used to call sSNV candidates. The regions covered by these read families corresponded to the actual callable regions in the Duplex-Multiome data, which are typically associated with open chromatin.

Since the sSNVs from the single-cell WGS data were called using the GRCh37 genome, we used the R package rtracklayer v1.48.0^59^ along with the liftOver chain file^60^ (GRCh38 to GRCh37) to lift over the callable regions. We then counted the number of sSNVs from the published single-cell WGS data within the Duplex-Multiome callable regions of the corresponding cell types and calculated the sSNV enrichment ratio in these callable regions as follows:

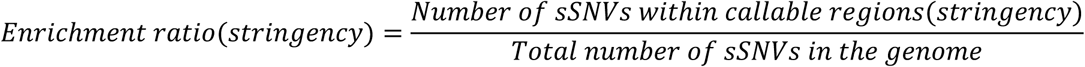

where *stringency* refers to the number of reads from each strand required for sSNV candidate calling.

### Sensitivity of variant detection for Duplex-Multiome

The bulk WGS BAM files were analyzed by HaplotypeCaller of GATK to identify germline variant candidates. Variants overlapping SNPs with a minor allele frequency of ≥1% in dbSNP v147 were extracted as high-quality germline variants. For each single cell, the high-quality germline variants within its callable region were counted (*G_all_*). Additionally, germline variants detected by Duplex-Multiome that matched the high-quality set were counted (*G_detected_*). The sensitivity for each cell was calculated as follows:

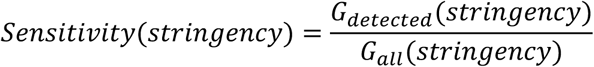

### Cell-type specific double-stranded sSNV burden estimation

A simulation-based maximum likelihood estimation method was developed to assess cell-type-specific double-stranded sSNV burdens. For each cell type or cluster of interest, variants detected within the cluster were counted to construct an observed probability distribution of variant counts per cell (*P_obs_*). Mutational burdens were modeled as discrete Gaussian distributions, based on prior scWGS data. To estimate the parameters (?) for the target cluster, a grid search was performed, during which mutation counts were sampled from the proposed distribution and mutations were randomly assigned across the combined callable regions of all cells in the cluster.

We noticed a small percentage of read families (<1%) displayed improper strand-barcoding. Specifically, each of our strand-barcoding steps utilized a pool of 4 different strand barcodes, resulting in a total of 4 unique strand1 barcodes and 4 unique strand2 barcodes. Properly strand-barcoded read families therefore contained 1 strand1 barcode and 1 strand2 barcode. However, we observed a small fraction of ‘multiple strand2 error’ read families containing 2 different types of strand2 barcodes. We realized that the design of our strand-tagging protocol meant that the fraction of these ‘multiple strand2 error’ read families would be equal to the fraction of ‘duplex error’ read families containing strand2 products improperly barcoded with 1 strand2 and 1 strand1 barcode. (Note that the fraction of ‘duplex error’ read families originating from strand1 products was negligible due to the design of our strand-tagging protocol, in which improperly barcoded strand1 products cannot be pre-amplified.) While ‘duplex error’ read families could not be directly distinguished from correctly barcoded read families, their fraction could be inferred from the fraction of ‘multiple strand2 error’ read families divided by 3/4, since the probability of a strand 2 randomly obtaining a strand 1 or 2 barcode is same and there is a probability of 1/4 in a ‘multiple strand2 error’ read family that strand2 products acquire the same strand2 barcode and therefore escape detection. The sSNV candidate rate in ‘duplex error’ read families was calculated using reads with only strand 2 barcodes, with the same parameters applied as in standard sSNV calling, except for the requirement of data from both strands. Next, we calculated the number of sSNV candidates contributed by the ‘duplex error’ read families (fraction of ‘duplex error’ read families * ratio of sSNV candidates within erroneous read families) and the number of simulated sSNV candidates from the correctly strand-barcoded read families. The sum of them times the sensitivity is the number of mutations we could detect in this round of simulation.

Simulations were run for at least 20,000 iterations to ensure convergence, and a distribution of variant counts per cell (*P_sim_*) was generated for the set of Ф. Likelihoods were calculated by comparing simulated and observed results under a Poisson distribution. Thethat maximized the likelihood was selected (Ф), and the corresponding sSNV burden for the target cell cluster was deduced.

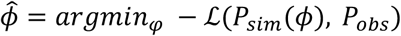

As described in the subsection “Calling of sSNV candidates from Duplex-Multiome data”, sSNV candidates identified under the “a6s3” stringency were used here. To minimize the risks associated with small sample sizes, this method was applied only to cell clusters containing more than 200 cells and with over 20,000,000 base pairs of callable regions to estimate mutational burden. If sufficient callable regions are obtained under the “a8s4” and/or “a10s5” stringency, this method will also be applied to sSNV candidates identified under these stricter criteria. In such cases, the estimated mutational burden for a cluster was determined as the minimum value calculated across all available stringencies. For clusters with extremely low mutational burden, where no sSNV candidates were identified under any stringency even “a2s1” despite having enough cells and length of callable regions, the estimated mutational burden was set to 0.

Using the enrichment ratios of variants between Duplex-Multiome callable regions and the whole genome, genome-wide somatic mutation burdens for excitatory neurons and oligodendrocytes were inferred. These estimates were consistent with previously published data.

### Calling of single-stranded DNA damage candidates from Duplex-Multiome data

We took advantage of the fact that each of our strand-tagging steps utilized 4 unique barcodes to perform higher accuracy same-strand consensus calling for single-stranded DNA damage candidates. Specifically, after the first round of amplification, strand 2 of a DNA fragment produced two copies. One of these obtains a strand 2 barcode through correct strand barcoding, while a small fraction of the other can randomly acquire another strand 2 barcode during amplification. These products will then be amplified and sequenced independently.

By comparing reads with different strand 2 barcodes from these read families, variants that appear only in reads with one barcode are likely artifacts introduced during amplification or sequencing. In contrast, variants present in reads with two different strand 2 barcodes could indicate real single-stranded DNA damage, double-stranded mutations (< 10%), or single-stranded lesions caused during sample storage or library construction.

### Cell-type specific single-stranded DNA damage burden estimation

Using these read families with two distinct strand 2 barcodes (*Callable region_ss_*), we identified single-stranded DNA damage events (3_))_), with the same parameters applied as in standard sSNV calling. We required that the variants be in reads labeled with two different strand barcodes.

The sensitivity (*Sensitivity_ss_*) was calculated as described in the subsection “Sensitivity of variant detection for Duplex-Multiome”, but only using these read families with two strand 2 barcodes.

Due to the limited number of read families with two strand 2 barcodes, our standard burden estimation method was not applicable in this context. Therefore, we directly calculated the single-stranded DNA damage burden as follows, though this approach does not account for standard deviations:

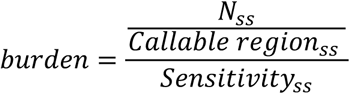

### Modeling aging effects on cell-type specific sSNV burden

Cell-type specific distributions of sSNV burden were calculated for different cell types of healthy brain samples. To estimate the aging effects on sSNV burden of cell type F, a value was randomly sampled from the distribution of this cell type for different individuals. Using these sampled values, a slope (*slope_ij_*) and an intercept (*intercept_ij_*) were calculated with the least squares method:

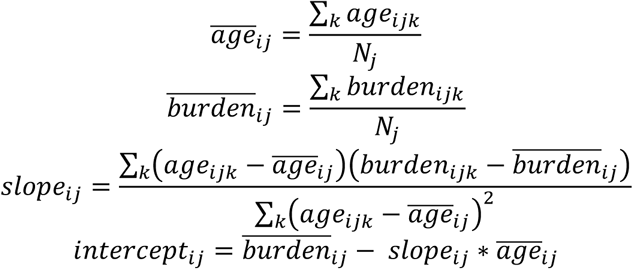

where *i* refers to the current round of sampling, *j* refers to the cell type, *k* refers to the individual, and *N_j_* is the number of samples where we have their cell type *j*’s sSNV burden available. The above steps were repeated for *M* iterations until convergence. Afterward, the slope, intercept, and standard deviations were calculated:

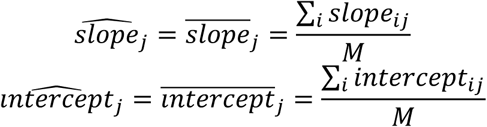

Statistical significance of the slope was next estimated based on one-sample *t*-test with *df* = *N_j_* − 2:

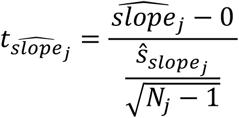

where *slope_j_* is the standard deviation of *slope_j_*

We compared the accumulation rates of sSNV burden across cell types. To determine if there is a statistically significant difference between a pair of cell types (cell type *a* and *b* as an example), we repeated the sampling procedure described earlier, but this time only retained the sampled values from *N_min_* individuals, where *N_min_* represents the smallest number of individuals with available sSNV burden data across cell types. This strategy was designed to alleviate the problem of sample size imbalance between the two cell types. We then applied a two-sample *t*-test:

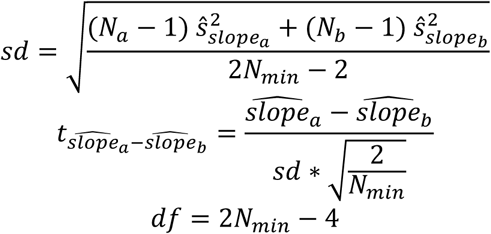

where *slope_a_* and *slope_b_* are the standard deviation of *slope_a_* and *slope_b_* respectively.

### Cell-type specific single-stranded DNA damage spectrum analysis

Single-stranded DNA damage candidates were combined for each cell type. The frequency of single-stranded DNA damage across the 96 trinucleotide contexts (*S_a_*) was analyzed using MutationalPatterns v3.0.1^61^. A previous study reported that the single-stranded DNA damage spectrum is similar to SBS30 from the COSMIC signatures^40^. To evaluate this, we compared the spectrum we obtained with SBS30 from COSMIC v3.2 and calculated cosine similarity scores for cell types with ≥ 200 cells from at least one individual.

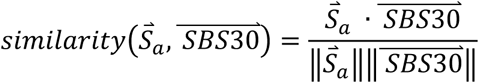

where *S_a_* refers to the single-stranded DNA damage spectrum of cell type *a*.

### Cell-type specific double-stranded sSNV spectrum analysis

Double-stranded sSNV candidates were combined for each cell type, and their frequency across the 96 trinucleotide contexts (*S^ds^_raw_*) were then calculated. To account for single-stranded DNA damage originating from “duplex error read families”, the single-stranded DNA damage spectrum for the corresponding cell type (calculated in the subsection “Cell-type specific single-stranded DNA damage spectrum analysis, *S^ds^_raw_*) were subtracted from the double-stranded sSNV candidate spectrum. This subtraction was weighted by 9, the fraction of sSNV candidates attributed to erroneous read families, as estimated in the subsection “Cell-type specific double-stranded sSNV burden estimation”.

To correct for bias in the background 96 trinucleotide context within Duplex-Multiome callable regions, the number of each trinucleotide type was counted within the callable regions of each cell type (*S^ds^_raw_*) and across the whole genome (*S^ds^_raw_*). The spectrum was then normalized using the following approach (similar to previous studies^2^):

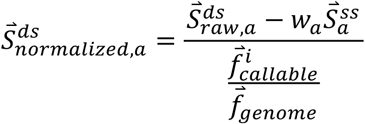

Here, + refers to the cell type of interest, and all division operations between two vectors were performed element-wise, which is:

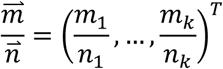

We compared the normalized spectra of excitatory neurons and oligodendrocytes with published data and calculated cosine similarity scores. Additionally, we calculated cosine similarity scores for all combinations of cell type pairs.

Given a previous study has identified five major mutational signatures (SBS1, SBS5, SBS16, SBS19, and SBS32) contributing to excitatory neurons and oligodendrocytes, we performed signature refitting with the five signatures on the normalized spectra of excitatory neurons and oligodendrocytes. This was done using non-negative least squares via the fit_to_signatures function in MutationalPatterns v3.0.1^61^.

### Duplex-Multiome covered region mutational signature enrichment analysis

As described in the subsection “Duplex-Multiome covered region sSNV enrichment analysis”, we downloaded the published single-cell WGS data for excitatory neurons and oligodendrocytes and lifted over our callable regions to match the reference genome, GRCh37. We calculated the frequency of sSNVs across the 96 trinucleotide contexts for these published single-cell WGS data (within the Duplex-Multiome callable regions and across the whole genome). The spectrum of sSNVs within the Duplex-Multiome callable regions was then normalized by the background 96 trinucleotide context for each cell type.

For a mutational signature from COSMIC, signature exposure levels were estimated for these spectra using the fit_to_signatures function in MutationalPatterns. The enrichment ratio was then calculated as follows:

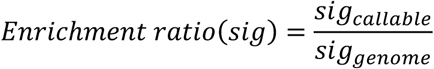

where *sig_callable_* and *sig_genome_* represent the signature exposure levels of a cell type in the Duplex-Multiome callable regions and the whole genome, respectively.

### Calling of clonal sSNV candidates from Duplex-Multiome data

Given the rarity of errors occurring at identical positions across cells, variants present in multiple cells but filtered out due to a low number of supporting reads are likely real. To enhance the detection of clonal sSNV candidates, we repeated the SNV calling pipeline and parameters described in the subsection “Calling of sSNV candidates from Duplex-Multiome data”, but we relaxed the criteria to require supporting reads ≥ 2 (denoted as a2s0) and allowed variants within read families shorter than 100 bps to be called.

Since our stringent germline mutation removal criteria could filter out sSNVs arising during early developmental stages, we adjusted the filtering approach for brain samples. In addition to variants that were not present in matched bulk WGS data, we retained those found in bulk WGS data but with a VAF < 10% as clonal sSNV candidates. For COLO829BLT-50, two cases were applied. 1) When using bulk WGS data from the COLO829BLT-50 cell line mixture to filter germline variants, the VAF threshold was lowered to < 40% to account for potentially larger clone sizes. 2) When using bulk WGS data from COLO829BL or COLO829T cells, the original threshold was applied, retaining only variants that were not present in matched bulk WGS data.

Due to the lowered thresholds, some real germline mutations could enter the clonal sSNV candidate pool. To address this, we implemented a germline variant filter. As described in the subsection “Sensitivity of variant detection for Duplex-Multiome”, variants overlapping SNPs with a minor allele frequency ≥ 1% in dbSNP^62^ were saved as high-quality germline variants. Next, we implemented a Bayesian linear regression model using Markov Chain Monte Carlo (MCMC) method (rstan v2.32.6^63^) to model the relationship between the number of cells with the high-quality germline mutations (M) and the total cells where we capture the region harboring the mutations (d) with Gaussian distribution for each sample. The model assumed:

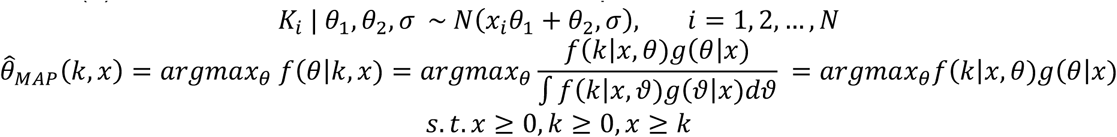

Theoretically, germline mutations should be detectable in approximately half of the cells capturing the region, as typically only one allele is captured at specific loci. Therefore, we assigned g_9_ with a prior 3(0.5, 100^/^), g_/_ with a prior 3(0, 10^/^), and h with a non-informative prior D+4%ℎ0(0, 10000). The MCMC sampling was performed using Hamiltonian Monte Carlo algorithm implemented in rstan v2.32.6. We used 4 chains with 2,000 iterations and a burn-in of 1,000 during modeling. Convergence was assessed through trace plots and Gelman-Rubin statistic R < 1.1. This generated a posterior predictive distribution of observing M cells with a germline mutation among d total cells where the mutation site was captured, ℎ(M, d,). This distribution was applied to identified clonal sSNV candidates and the probability of being a germline mutation could be derived. This probability was then applied to filter potential germline mutations from the clonal sSNV candidate pool. To avoid the multiple testing problem, only clonal sSNV candidates with

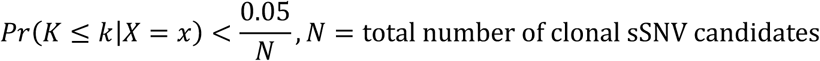

were retained for the downstream analysis. In this work, we had 332 clonal sSNV candidates for healthy brain samples, so we used *P* < 0.00001 < 0.05 as the threshold.

Systematic mapping errors at specific genomic locations can lead to the same false-positive variants being identified across multiple cells, incorrectly classifying them as clonal sSNV candidates. Local realignment is the common way to avoid mapping errors. Lines corresponding to the read families containing the identified clonal sSNV candidates were extracted from the BAM files of the single cells. These extracted BAM files were realigned using BWA, with a reduced gap open penalty of 4, which allows the aligner to better recognize Indels. The realigned BAM files were then used to re-identify variants using the same parameters as described earlier. In addition, the extracted BAM files were analyzed by HaplotypeCaller of GATK, which has more layers of filtering including local realignment. Variants that were identified by both approaches in at least 50% of the cells were considered robust and saved as clonal sSNVs.

### Recovery of cells with clonal sSNVs and low-quality transcriptome data

Cells with low-quality transcriptome data were excluded during the filtering process (see details in the subsection “10x single nucleus multiome data analysis”), so their cell types were undetermined. For cells in which clonal sSNVs were detected but transcriptome data quality was not good enough, we attempted to recover them if their ATAC-seq data met quality standards. We applied the same QC criterion as previously described (nCount_ATAC > 1000) and utilized the pipeline detailed in the subsection “10x single nucleus multiome data analysis”. To process the snATAC-seq data, we merged data from all subjects and conducted RunTFIDF, FindTopFeatures, and RunSVD. Batch effects were corrected using Harmony. The integrated snATAC-seq data was further analyzed using FindNeighbors, RunUMAP, and FindClusters.

For cells with high-quality RNA and ATAC data, cell type had been annotated based on marker gene expression. The predominant cell type in each cluster generated from the snATAC-seq data was assigned as the cluster identity.

### Inferring lineage relationships via shared sSNV graph

Graph-based techniques were applied to analyze clonality based on clonal sSNVs. Cells sharing the same clonal sSNVs were connected, and the distance between each cell pair was calculated based on the number of shared clonal sSNVs. This analysis, which was implemented using the R package igraph v1.2.5^64^, generated a complex graph structure representing cell lineage relationships. To identify distinct clones, we applied an unsupervised machine learning algorithm, label propagation, to the graph. The resulting structure was projected onto a 2D plane for visualization.

### Clonal sSNV spectrum analysis

For COLO829BLT-50, the frequency of clonal sSNVs was calculated across the 96 trinucleotide contexts separately for these passed filtering using bulk WGS data from either the COLO829BLT-50 cell line mixture, or COLO829BL cells, or COLO829T cells.

For clonal sSNVs from control brain samples, the frequency of sSNVs across the seven types of base substitutions (distinguishing C>T substitutions at CpG sites from non-CpG sites) were analyzed by mut_type_occurrences of MutationalPatterns. Also, for each individual, the frequency of clonal sSNVs across the 96 trinucleotide contexts was calculated. To address bias in the background 96 trinucleotide context, the callable regions for clonal sSNVs were identified for each individual. Since the clonal sSNV calling pipeline requires sufficient depth at each locus, the minimum number of cells in which the identified clonal sSNV positions were captured for each individual represents the required depth for the method to reliably detect clonal variants in that individual. Hence, regions that were covered in more than the minimum number of cells were the actual callable regions for the individual. For individuals without identified clonal sSNVs, the average callable region size from other individuals was used as an estimate. The frequency of each trinucleotide type was then counted within the callable regions for each individual and across the whole genome. Finally, the clonal sSNV spectrum was normalized based on the background trinucleotide context and aggregated to generate the normalized clonal sSNV spectrum.

We performed signature analysis with the COSMIC mutational signatures on these spectra, which was done using non-negative least squares via the fit_to_signatures function in MutationalPatterns^61^.

### Clonal sSNV rate analysis

Cell-type-specific clonal sSNV rates were calculated using a similar approach. For each cell type in a sample, we counted the total number of cells harboring each clonal sSNV. To estimate the callable region size for each cell type, we summed the covered lengths of callable regions from all single cells within the cell type. The clonal sSNV rate for each cell type in a sample was then calculated as 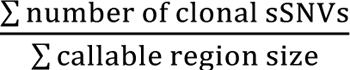. We next pooled cell types for each sample to estimate clonal sSNV rates for all samples. A linear regression was then conducted to determine whether any samples showed significantly higher or lower clonal sSNV rates. To compare the rates between neurons and glial cells, we conducted a paired one-tailed Wilcoxon test.

### Amplicon sequencing validation for the identified clonal sSNVs

Amplicons from 200-500bp centered around clonal variants were produced via PCR from bulk DNA extracted from the same brain tissue Duplex-Multiome was performed on and sequenced on the MiSeq via the Amplicon-EZ service from Azenta. Sequencing reads were aligned to the GRCh38 reference genome using BWA and sorted using the SortSam function in Picard. The resulting BAM files were further processed with GATK v3.6 for indel realignment to enhance the accuracy of mutation detection.

For each somatic mutation candidate, MosaicHunter v1.0 filtered out the low-quality reads and calculated the number of reads supporting each allele. To confirm a candidate as a somatic mutation, the read fraction of the mutant allele needed to exceed the sum of the fractions of the next two most common error alleles. The results were then manually inspected using IGV^65^.

### Genotyping known mutations in single cells from Duplex-Multiome data

For clonal variants identified through deeply sequenced bulk WGS data (e.g., variant chr1-205284910-T-C from sample AN06365), genotyping was performed for individual cells from the same sample using BAM files generated in the subsection “Calling of sSNV candidates from Duplex-Multiome data”. The following criteria were used to identify the variant: base quality > 30, MapQ > 20, supporting reads ≥ 2, and a read family VAF of 1. Cells with this variant detected were saved for the downstream analysis. To ensure accuracy, all results were manually validated using the Integrative Genomics Viewer (IGV) v2.16.0^65^.

### Association analysis of clonal sSNVs and surrounding gene expression changes

We developed an empirical statistical model using bootstrapping to explore the mutational effects on gene expression, addressing the limitation that Duplex-Multiome may not detect mutations in all cells harboring them due to allelic dropout. For each clonal sSNV, a pseudobulk expression profile was created from the cells in which the variant was detected. This profile included genes located within 500,000 base pairs on either side of the variant (e.g., M genes as an example).

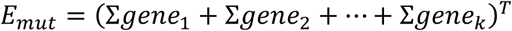

Next, we generated pseudo-clone data by sampling an equivalent number of cells as in the previous step from the same cell type within the same individual, excluding the cells where the variant was detected. For each pseudo-clone, a pseudobulk expression profile was produced, containing the same genes as in the original pseudobulk profile. Through *n* rounds of permutation (with *n* chosen to ensure that each cell is sampled at least once), we obtained enough bootstrapped data. These bootstrapped profiles were then used to construct empirical distributions of gene expression across the pseudo-clones.

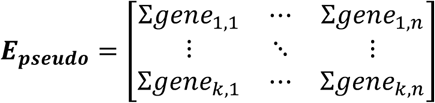

Here, the columns represented pseudo-clones, and the rows corresponded to genes located near the variant of interest. To compare the pseudobulk expression profile of mutant cells with these empirical distributions, we ranked the expression levels of each gene in the mutant cell profile relative to the bootstrapped data. This ranking provided a statistical significance value for each mutation-gene pair, indicating how the presence of the mutation influenced the expression of nearby genes.

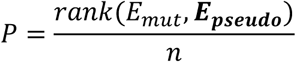

### Luciferase assays

Site-directed mutagenesis (New England Biolabs) and cloning was carried out on one identified sSNV mutation to recreate the wildtype and mutant constructs. Constructs were cloned into the luciferase vector pNL3.2[NlucP/minP] (Promega), and cloning was successful for both the wildtype and mutant. The luciferase constructs were then transfected into N2A cells along with the normalizer vector pGL4.54[luc/TK] (Promega) with Lipofectamine 3000 (Thermo Fisher).

Three technical replicates were done per biological replicate across a total of four biological replicates. Luciferase assays were then carried out 24 hours post transfection using the Nano-Glo® Dual-Luciferase® Reporter Assay System (Promega) with values averaged across technical replicates.

## Data Availability

Data will be deposited in dbGAP or NIAGADS and accession codes made available before publication.

## Code Availability

The source code for Duplex-Multiome analysis pipeline and other scripts for analyses described in this study are available on GitHub (https://github.com/ShulinMao/Duplex-Multiome/tree/dev_v1)

## Acknowledgements

We thank the NIH Neurobiobank at the University of Maryland Brain and Tissue Bank, Boston University UNITE or VA-BU-CLF Brain Bank and the Autism Brain Net for sharing their tissue collections, and we are grateful to the donors and their families for their invaluable contribution to the advancement of science. We thank R. Mathieu at the Boston Children’s Hospital and Harvard Stem Cell Institute Flow Cytometry Research Facility, R. S. Hill, the Research Computing group at Harvard Medical School and the Boston Children’s Hospital Intellectual and Developmental Disabilities Research Center (IDDRC) Molecular Genetics Core for assistance. We thank the Somatic Mosaicism across Human Tissues (SMaHT) UW-SCRI Genome Characterization Center for generating and providing the COLO829 cell line mixture (BLT-50). This work was supported by National Institute of Health (NIH) grants R01AG070921 (C.A.W. and E.A.L.), R01NS032457 (C.A.W.), R01AG078929 (E.A.L and C.A.W.), R56AG079857 (A.Y.H., E.A.L., and C.A.W.), DP2AG072437 (E.A.L), R01AG088082 (A.Y.H.), and T32GM007748 (R.E.A.); John Templeton Foundation grant 62587 (the opinions expressed in this publication are those of the authors and do not necessarily reflect the views of the John Templeton Foundation) (C.A.W.); Simons Foundation grant 953759 (C.A.W.); Suh Kyungbae Foundation (E.A.L.); Allen Discovery Center for Human Brain Evolution, a Paul G. Allen Frontiers Group advised program of the Paul G. Allen Family Foundation (C.A.W. and E.A.L); Alzheimer’s Association Research Fellowship AARF-22-972287 (A.Y.H.); HHMI Jane Coffin Childs Fellowship (A.J.K.); American Heart Association Predoctoral Fellowship (S.M.); and Autism Speaks Postdoctoral Fellowship 13008 (R.E.A.). C.A.W. is an Investigator of the Howard Hughes Medical Institute. Fig. 1A-B, 2A, 3A and C, 5A, and 6A were created in https://BioRender.com.

## Author Contributions

A.J.K. and S.M. conceived the study; A.J.K. performed nuclear sorting and Duplex-Multiome; S.M. led computational analyses; A.J.K., D.D.S., and R.E.A. developed the Duplex-Multiome experimental method; D.A.S. developed DuplexTools; G.D. contributed to PTA analysis and C.C.M. contributed to PTA experiments; H.C. performed amplicon-sequencing and luciferase assays; A.Y.H., E.A.L. and C.A.W. directed the research and edited the manuscript; A.J.K. and S.M. wrote the manuscript.

---

**Extended Data Fig. 1.**
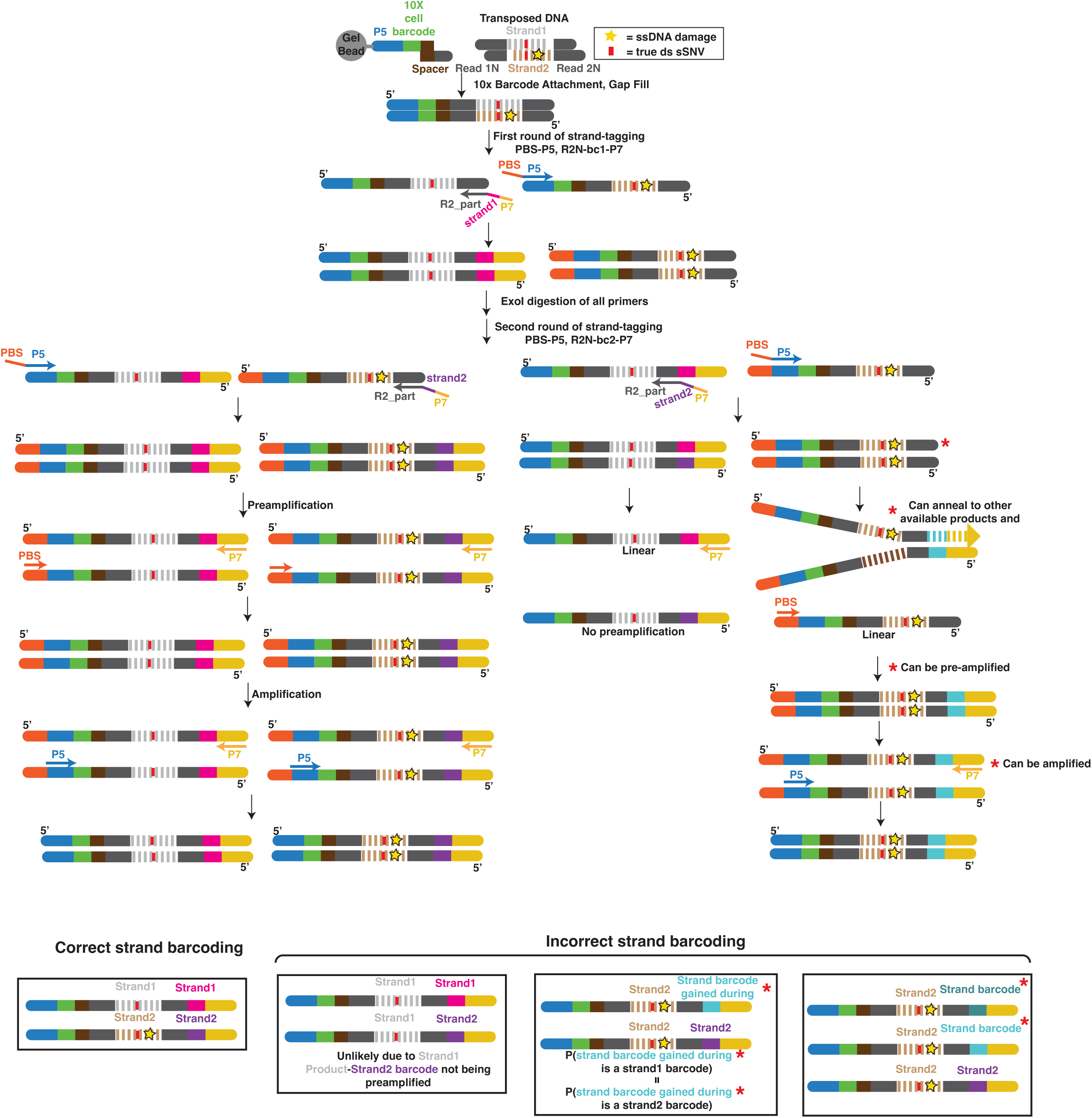
Full schematic of the Duplex-Multiome method. Detailed workflow of Duplex-Multiome starting from the GEM Generation & Barcoding step of the 10X Multiome protocol and tracing all products generated from both strands of an initial starting duplex of DNA. Steps not shown are identical to the published manual. Single-stranded DNA damage (yellow star) and true double-stranded DNA single-nucleotide variants (red rectangle) are tracked throughout. Correctly strand-barcoded products are illustrated in addition to read families generated after rare strand barcoding errors, as well as relative probabilities of each (Methods).

**Extended Data Fig. 2.**
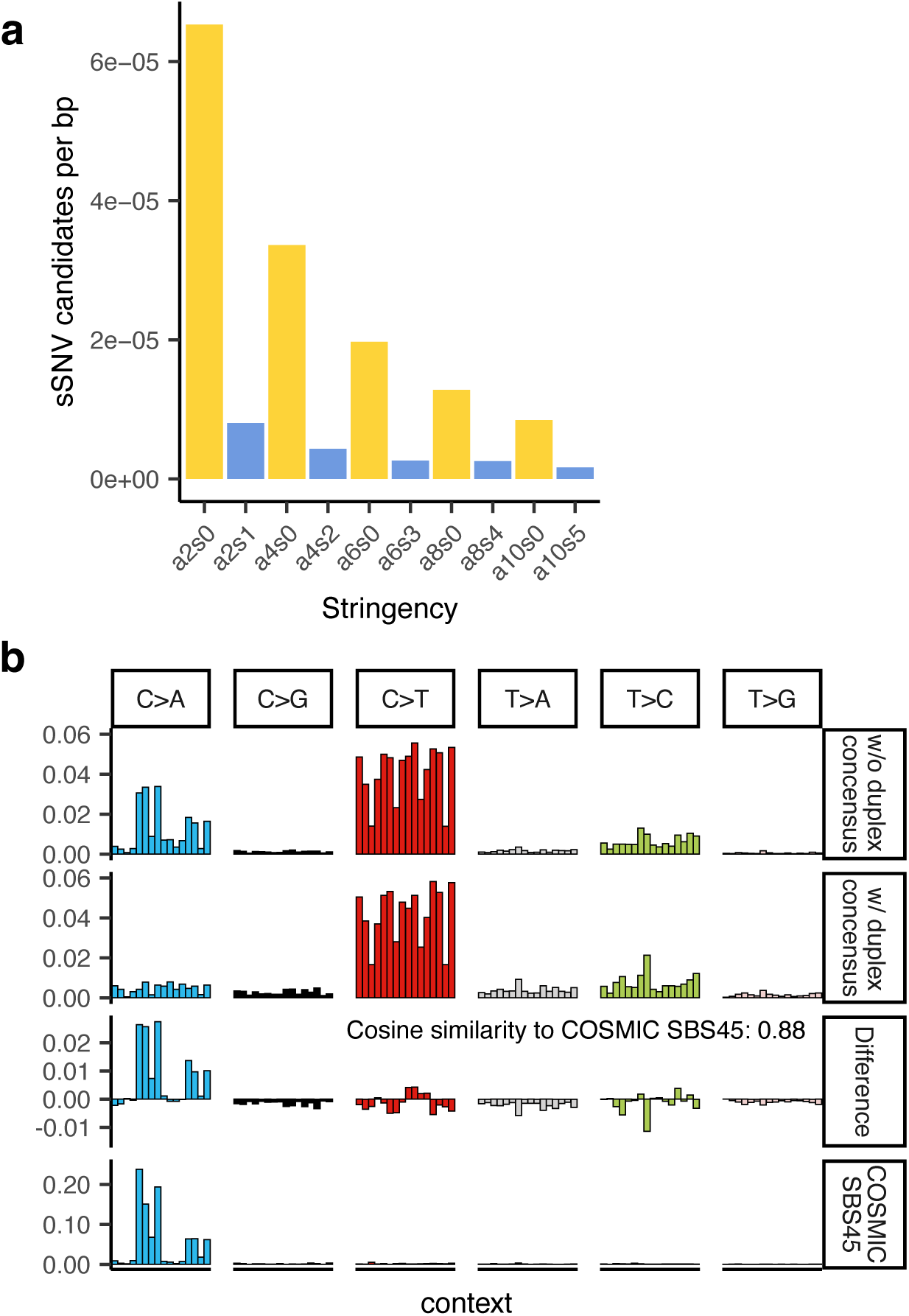
Duplex consensus removes artifacts and enables high-quality variant calling for Duplex-Multiome. **a.** sSNV candidates per base pair in UMB1932 under different stringency levels. Yellow bars indicate no requirement of duplex consensus, while blue bars require duplex consensus. The notations “a2s0”, “a4s0”, “a6s0”, “a8s0”, and “a10s0” represent stringency levels that require at least 2, 4, 6, 8, and 10 supporting reads, respectively, to call a variant but do not require duplex consensus. “a2s1”, “a4s2”, “a6s3”, “a8s4”, and “a10s5” denote stringency levels that require at least 1, 2, 3, 4, and 5 supporting reads from each strand to call a variant, which means duplex consensus is required. **b.** Mutational spectra of pooled sSNVs for stringencies “a2s0” (no duplex consensus) and “a2s1” (duplex consensus) from UMB1932. The difference spectrum and SBS45 signature that is associated with 8-oxo-guanine introduced during sequencing are also shown. Cosine similarity between the difference spectrum and SBS45 is calculated (Methods).

**Extended Data Fig. 3.**
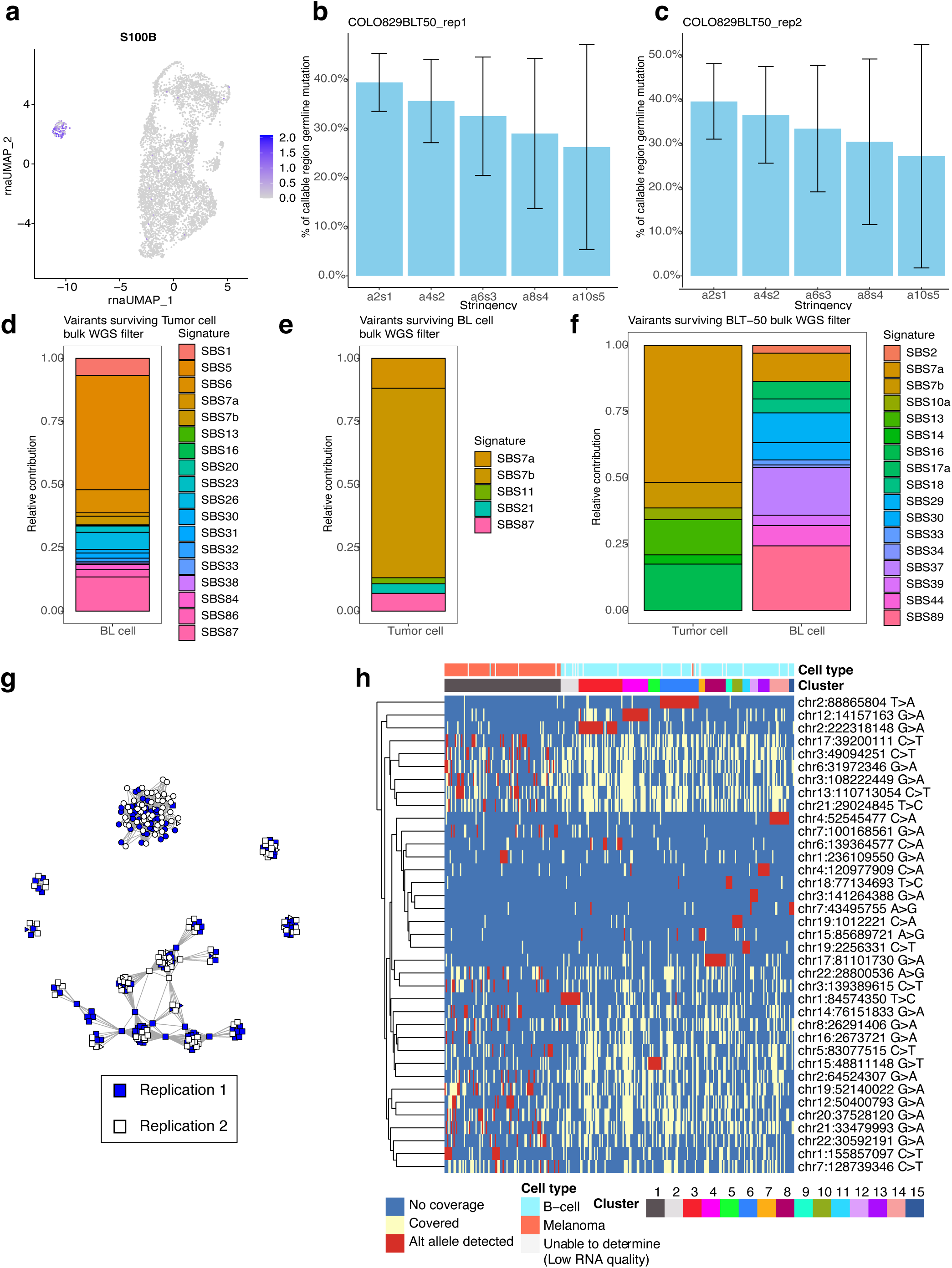
Duplex-Multiome profiles cell-type-specific mutational patterns and lineage relationships in COLO829BLT-50. **a.** UMAP of COLO829T marker *S100B* expression, identifying melanoma cells. **b – c.** Sensitivity of variant detection for Duplex-Multiome under different stringencies across two technical replications. The panels show that there are no significant batch effects between replications. Error bars indicate ±1 standard deviation across cells. **d - f.** Contributions of COSMIC SBS signatures to clonal sSNVs identified in tumor and BL cells, demonstrating differences in mutational processes between the two populations. **d.** Signature contributions for clonal SNVs in bulk BL cells filtered using bulk WGS from tumor cells. **e.** Contributions for tumor cell clonal SNVs filtered using bulk WGS from BL cells. **f.** Signature contributions of SNVs identified under the “a6s3” condition for both BL and tumor cells. **g.** Network graph of clonal sSNVs across two technical replicates. Nodes represent single cells with detected clonal sSNVs, and edges indicate the reciprocal number of shared variants (Methods). Cells are colored based on which replication they are from. Cells from the two replications are evenly mixed in the graph, suggesting there are no significant batch effects. **h.** Heatmap of clonal SNVs used to infer lineage relationships. Rows represent individual clonal SNVs, and columns show cells with clonal SNVs detected. Blue indicates no coverage of the site, light yellow indicates coverage without detection, and red indicates detection of the SNV. Clusters are annotated with cell types based on marker expressions (Methods). Cells with insufficient RNA quality are labeled as “unable to determine”.

**Extended Data Fig. 4.**
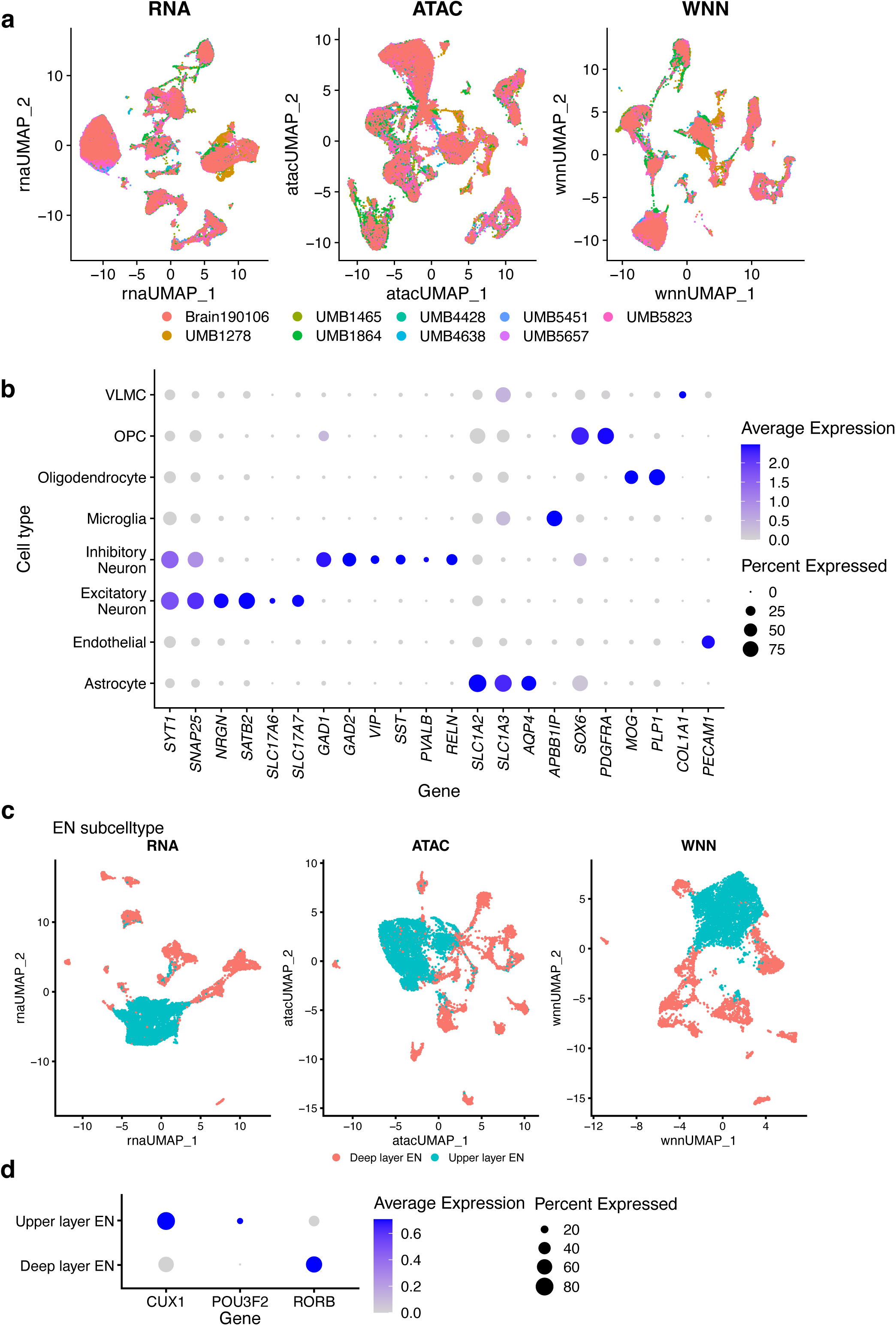
Cell type and subtype annotation in the aging human brain. **a.** UMAP plots of snRNA-seq (left), snATAC-seq (middle), and integrated snRNA-seq and snATAC-seq data using WNN analysis (right) from nine subjects. Each plot is colored by the subject of origin. **b.** Dot plot showing marker gene expression across annotated brain cell types. Dot size represents the percentage of cells in which the marker is detected, while color represents the average expression level in those cells. **c.** Cells that were annotated as ENs were selected for subtype annotation (Methods). We generated a separate Seurat object for these ENs. This panel shows the ENs’ UMAP plot snRNA-seq (left), snATAC-seq (middle), and integrated snRNA-seq and snATAC-seq data using WNN analysis (right) from nine subjects. Based on markers of EN subtypes, we annotated and labelled them in this graph. **d.** Dot plot showing expression of EN subtype markers. Legend details are consistent with panel **b**.

**Extended Data Fig. 5.**
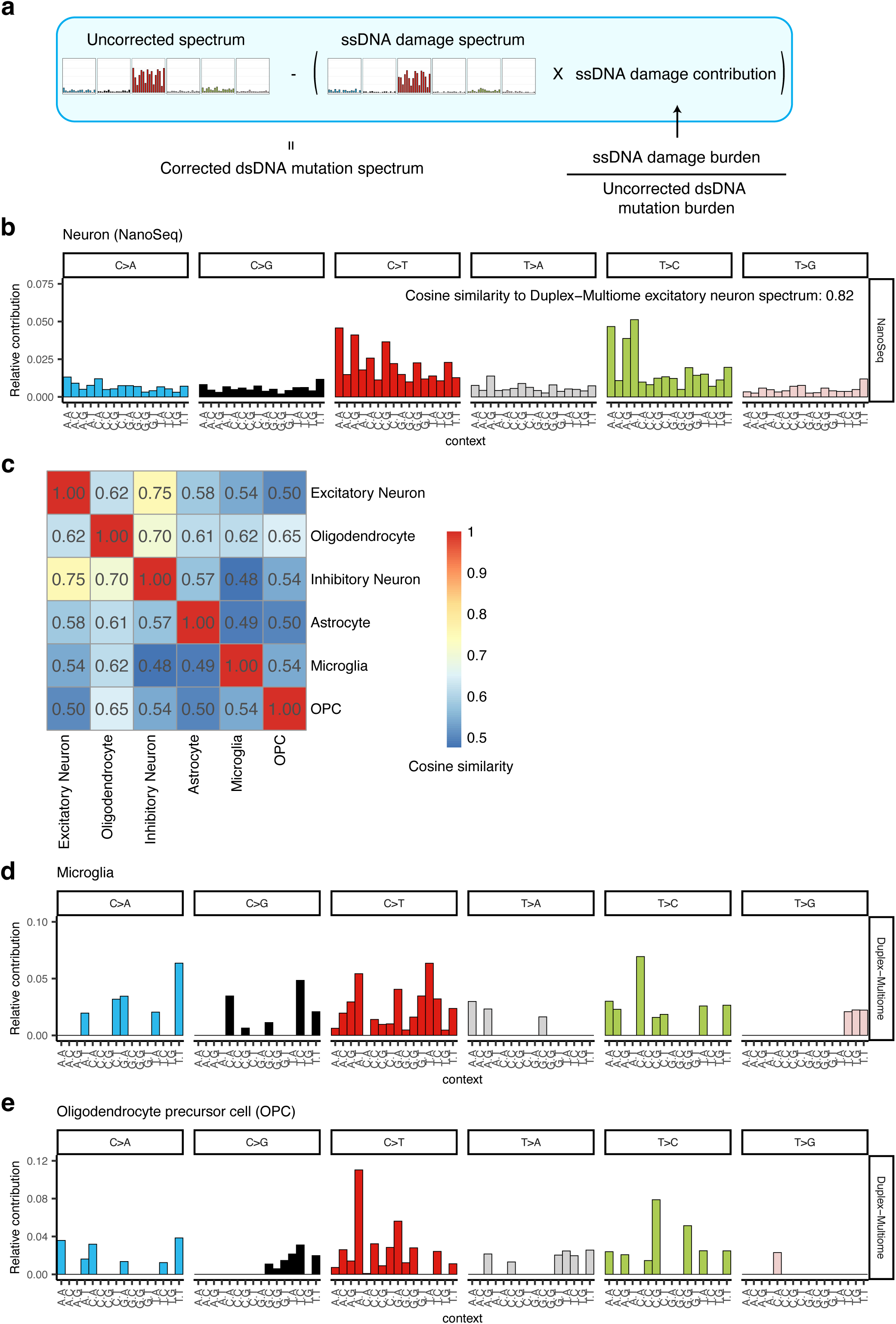
Mutational spectrum analysis of sSNVs in the aging human brain. **a.** Computational pipeline for correcting spectra of sSNVs. Uncorrected spectra were generated from raw sSNV candidates called by Duplex-Multiome, and ssDNA damage spectra were generated from ssDNA calls from Duplex-Multiome. Corrected spectra are obtained by subtracting ssDNA contributions from uncorrected spectra (Methods). **b.** Mutational spectrum of sSNVs identified from neurons by NanoSeq. The cosine similarity between it and the spectrum of ENs by Duplex-Multiome is shown. **c.** Heatmap of cosine similarity between sSNV spectra of different cell type pairs. The spectra of ENs and INs show the greatest similarity. **d - e**. Mutational spectra of pooled Microglia (panel **d**) and OPC (panel **e**) sSNVs identified by Duplex-Multiome.

**Extended Data Fig. 6.**
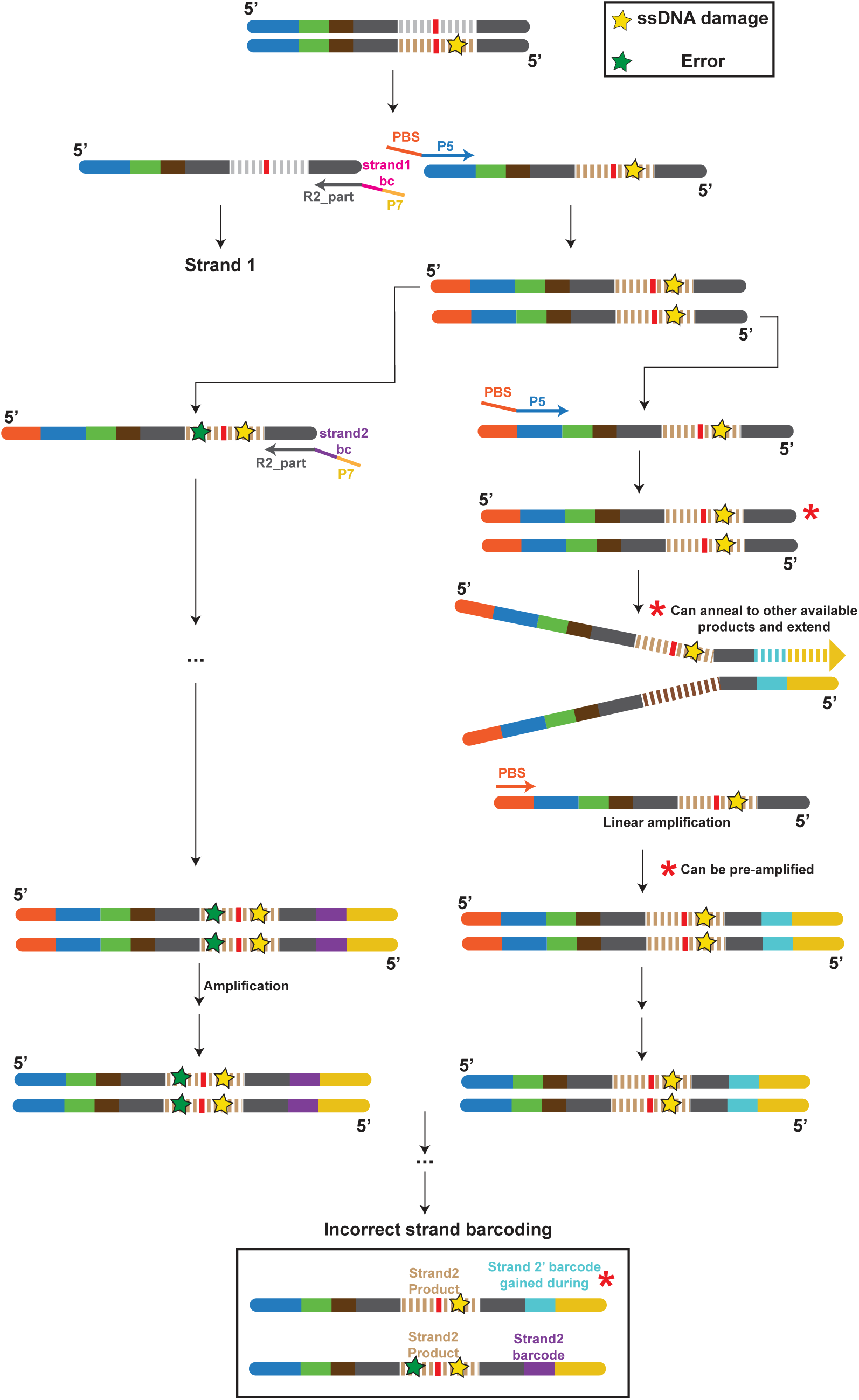
Duplex-Multiome strand-tagging approach enables same-strand duplex consensus. A schematic showing how strand-specific barcoding after the first round of amplification allows for same-strand duplex consensus. This approach distinguishes single-stranded DNA (ssDNA) damage from sequencing and amplification errors that occur after the first round (Methods).

**Extended Data Fig. 7.**
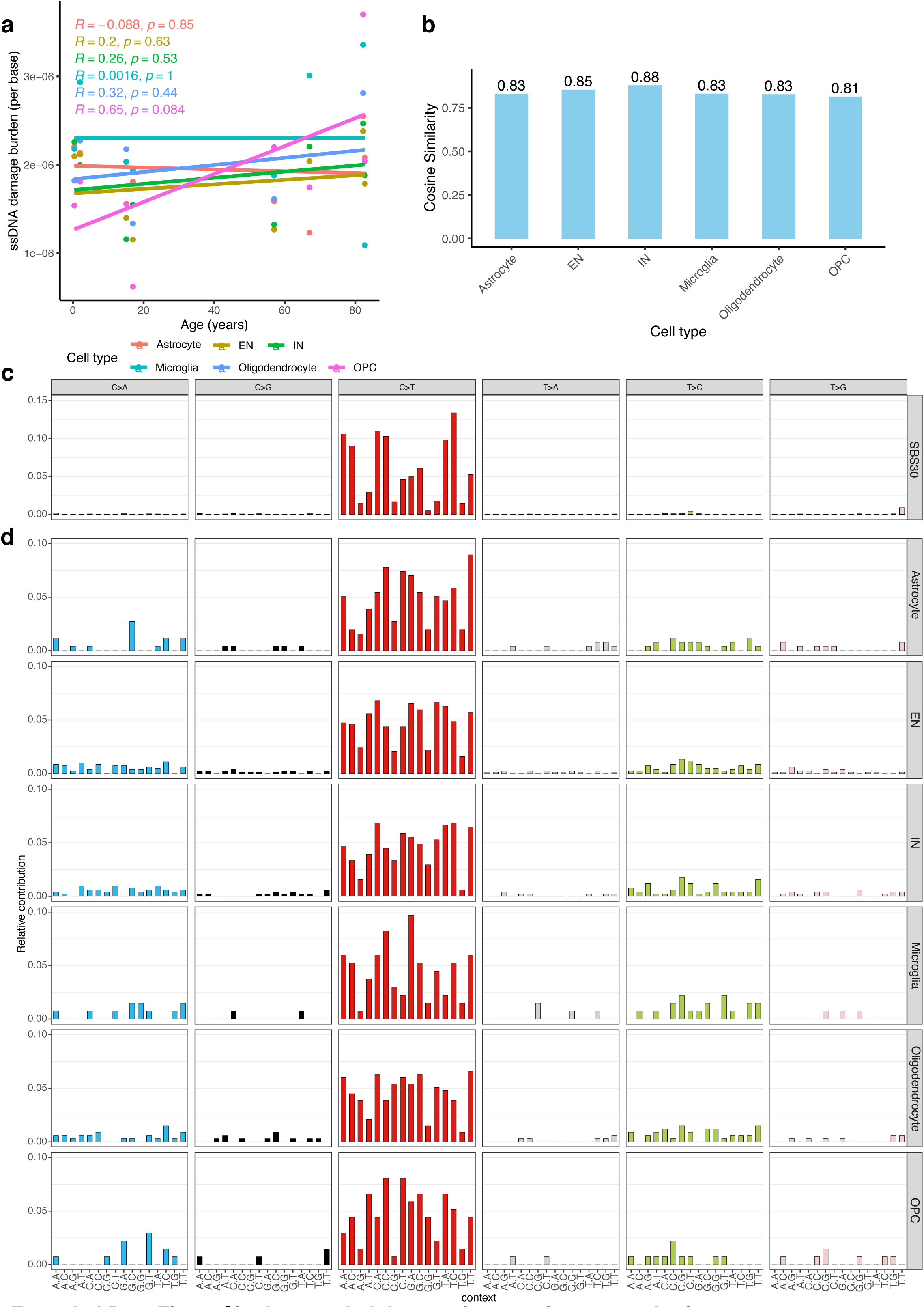
Single-stranded damage in the aging human brain. **a.** Cell-type-specific ssDNA damage burden in the aging brain. Each data point represents a cell type from a sample, colored by cell type. Linear trend lines are fitted to each cell type, with *R* and *P* values shown. **b.** Cosine similarity of ssDNA damage spectra for each cell type compared to COSMIC signature SBS30 that could reflect ssDNA damage spectrum. **c.** Mutational spectrum of SBS30. **d.** Mutational spectra of ssDNA damage for each cell type.

**Extended Data Fig. 8.**
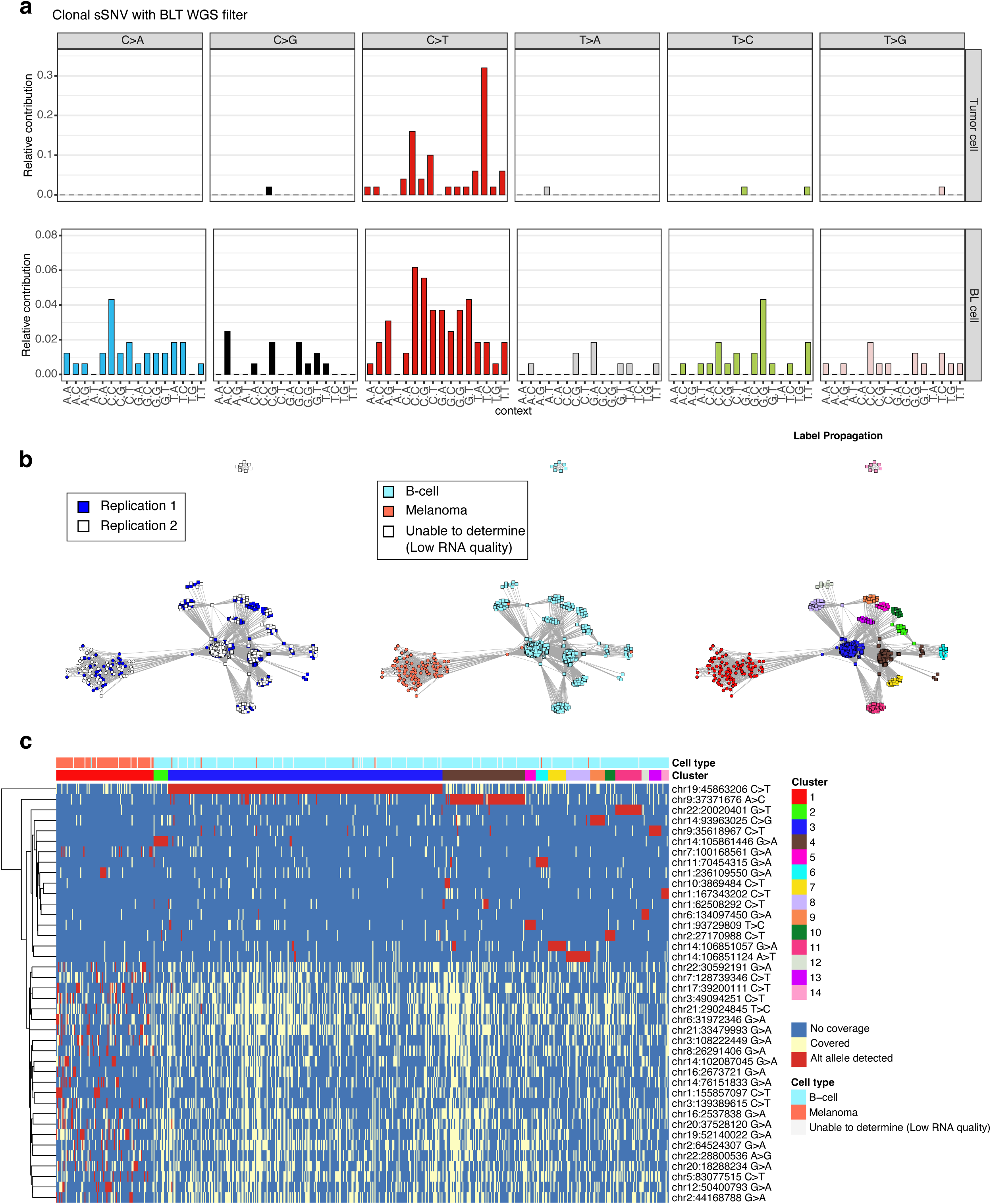
Benchmarking the clonal mutation analysis approach in COLO829BLT-50. **a.** Mutational spectrum of clonal sSNVs in tumor (upper row) and BL cells (bottom row) called using the clonal mutation calling pipeline (Methods). **b.** Inferred lineage relationships using clonal sSNVs in the panel **a**. The left panel shows two replications; the middle panel shows cell types annotated by marker expressions; the right panel represents the clusters annotated by label propagation (Methods). **c.** Heatmap of the clonal SNVs. Rows represent individual clonal SNVs, and columns show cells with clonal SNVs detected. Blue indicates no coverage of the site, light yellow indicates coverage without detection, and red indicates detection of the SNV. Clusters are annotated with cell types based on marker expressions (Methods). Cells with insufficient RNA quality are labeled as “unable to determine”.

**Extended Data Fig. 9.**
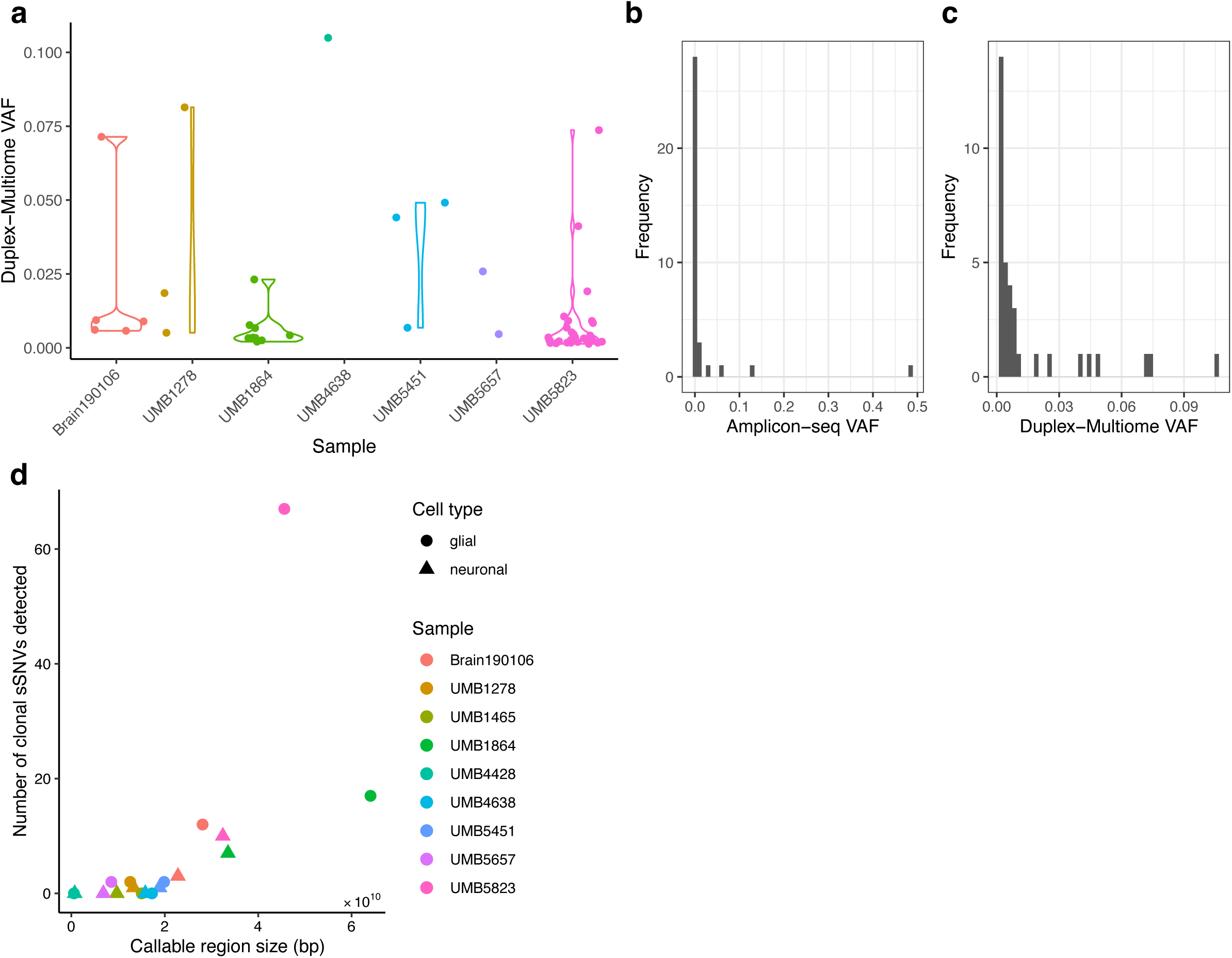
Cell-type-specific clonal sSNV rates and validation by Amplicon-seq. **a.** Violin plot showing Duplex-Multiome VAF of clonal sSNVs in each sample. **b.** Distribution of Amplicon-seq VAF for validated clonal sSNVs. **c.** Distribution of Duplex-Multiome VAF for validated clonal sSNVs. **d.** Number of clonal sSNVs detected in glial cells and neurons from each sample. The x-axis represents the corresponding callable region size. Round dots indicate glial cells from individual samples, while triangles represent neurons. All dots are colored by samples.

**Extended Data Fig. 10.**
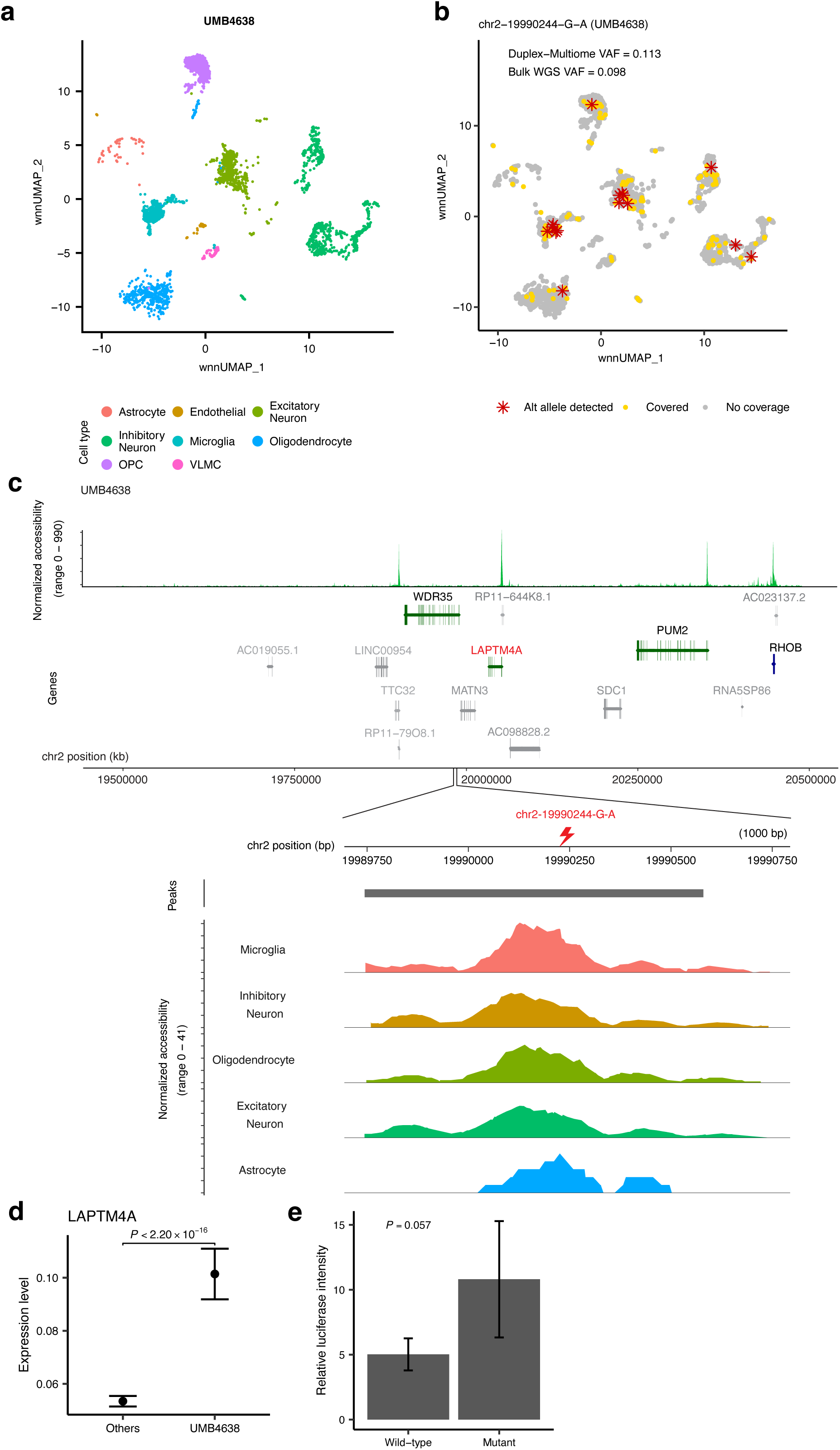
A neurotypical sample harbors a non-coding somatic mutation (chr2:19990244 G>A) correlating with changes in nearby gene expression. **a.** UMAP plot of integrated snRNA-seq and snATAC-seq data using WNN analysis for UMB4638. **b.** UMAP plot showing the G>A variant at chr2:19990244 from UMB4638. Duplex-Multiome detected the variant with a VAF of 0.113 and the previous bulk WGS detected it with a VAF of 0.098. Red stars indicate cells where the variant (alt allele) was detected by Duplex-Multiome. Yellow dots represent cells where the variant site was covered by Duplex-Multiome (either alt or ref allele could be detected). Grey dots denote cells where the variant site was not covered by Duplex-Multiome. **c.** Analysis of the G>A variant as in Figure 6a. Genes within a 1Mb interval of the variant are shown, with those expressed in fewer than 10% of cells excluded (colored grey). Blue and green indicate transcription direction of analyzed genes. Peaks from snATAC-seq data are displayed. The first and the bottom panel shows normalized snATAC-seq signals across the region. **d.** *LAPTM4A* expression in UMB4638 compared to other neurotypical brains. Data points represent mean expression values, with error bars indicating the 95% confidence interval. *P* is calculated with a two-sided Wilcoxon test. **e.** A construct containing the G>A variant at chr2:19990244 transfected into N2A cells had ∼2-fold higher promoter activity compared to a reference construct via luciferase assay (n=4 independent plates, *P* is calculated with a two-sided Wilcoxon test.).

