## Supplementary Figures for "Cell-type-specific patterns and consequences of somatic mutation in development and aging brain"

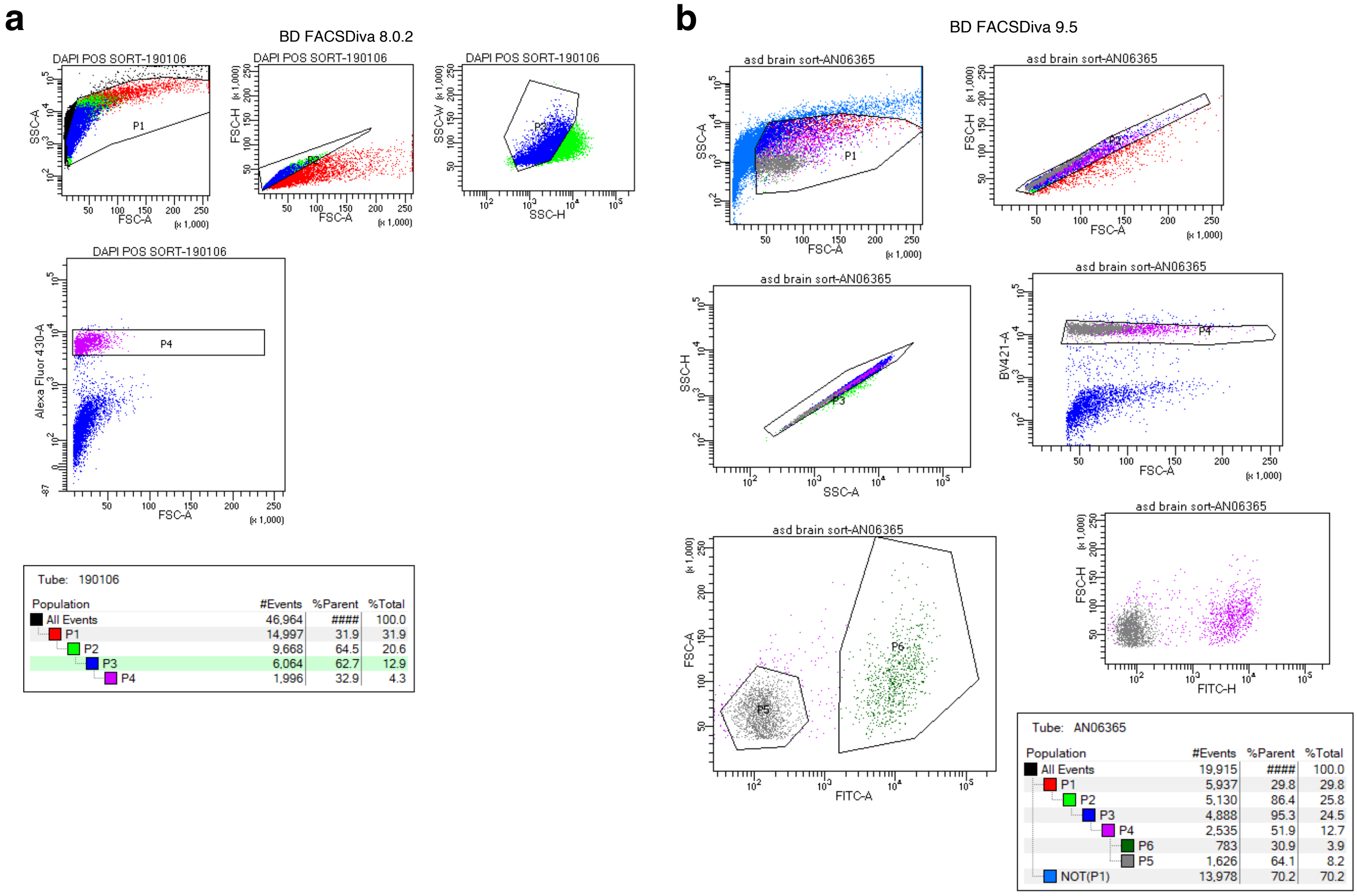


Tube: 190106

| Population | #Events | %Parent | %Total |
|---|---|---|---|
| All Events | 46,964 | #### | 100.0 |
| P1 | 14,997 | 31.9 | 31.9 |
| P2 | 9,668 | 64.5 | 20.6 |
| P3 | 6,064 | 62.7 | 12.9 |
| P4 | 1,996 | 32.9 | 4.3 |

Tube: AN06365

| Population | #Events | %Parent | %Total |
|---|---|---|---|
| All Events | 19,915 | #### | 100.0 |
| P1 | 5,937 | 29.8 | 29.8 |
| P2 | 5,130 | 86.4 | 25.8 |
| P3 | 4,888 | 95.3 | 24.5 |
| P4 | 2,535 | 51.9 | 12.7 |
| P6 | 783 | 30.9 | 3.9 |
| P5 | 1,626 | 64.1 | 8.2 |
| NOT(P1) | 13,978 | 70.2 | 70.2 |

**Supplementary Fig 1. FANS gating strategy for Dapi+ and NeuN+ sorting.**

**Supplementary Fig. 1. FANS gating strategy for Dapi+ and NeuN+ sorting.**

**a.** Representative FANS using Dapi to label nuclei for separation from debris. Box labeled ‘P4’ denotes events sorted as Dapi+ nuclei.

**b.** Representative FANS using AF488-conjugated anti-NeuN antibodies to label neurons and non-neuronal cells. Box labeled ‘P4’ denotes events sorted as Dapi+ nuclei, box labeled ‘P5’ denotes Dapi+ nuclei sorted as NeuN-negative, non-neuronal cells. Box labeled ‘P6’ denotes Dapi+ nuclei sorted as NeuN-positive, neuronal cells.

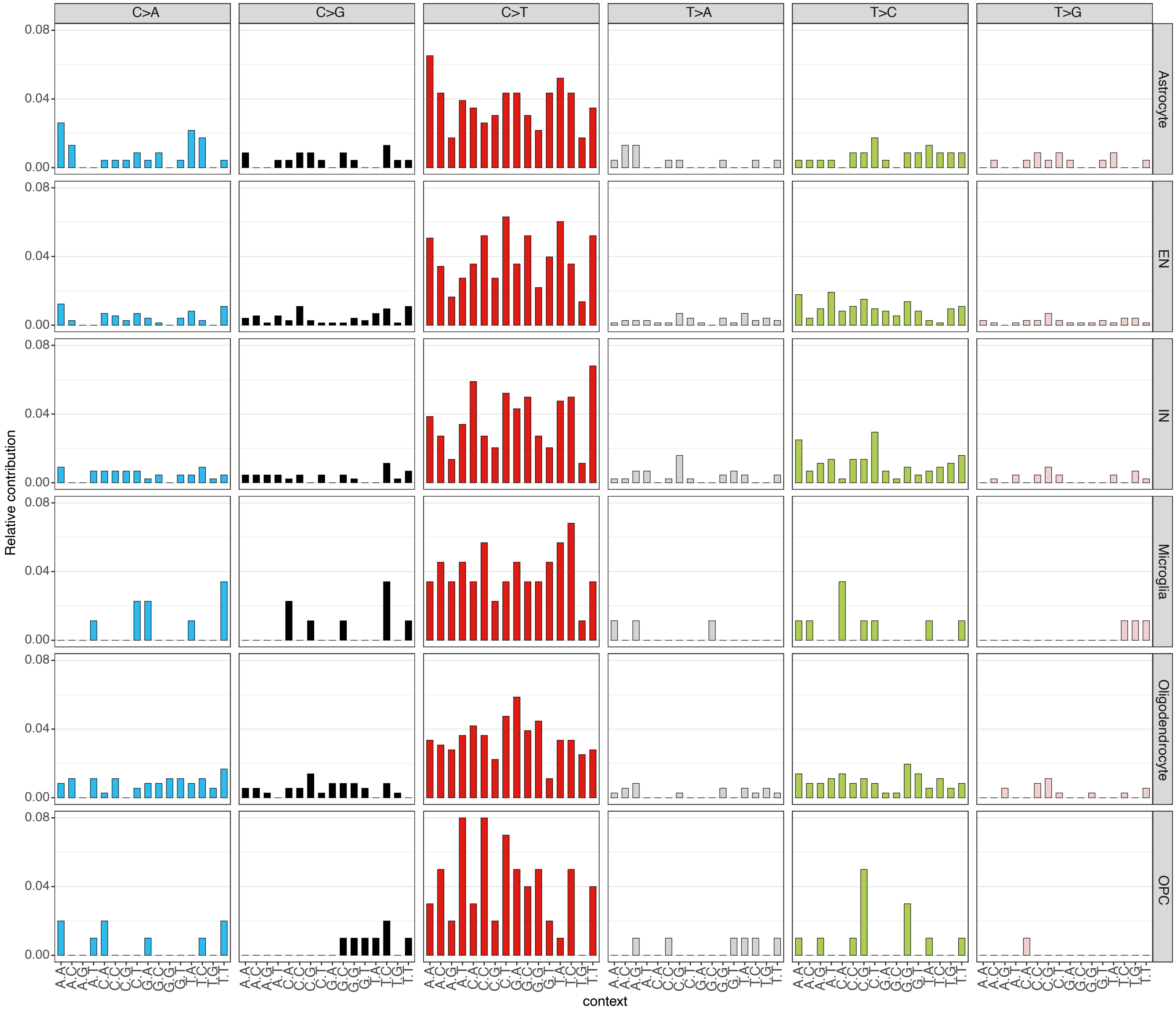


**Supplementary Fig. 2. Uncorrected spectra of raw sSNV candidates called by Duplex-Multiome.**

**Supplementary Fig. 2. Uncorrected spectra of raw sSNV candidates called by Duplex-Multiome.**

Mutational spectra of pooled raw sSNV candidates (under a6s3) for each cell type.

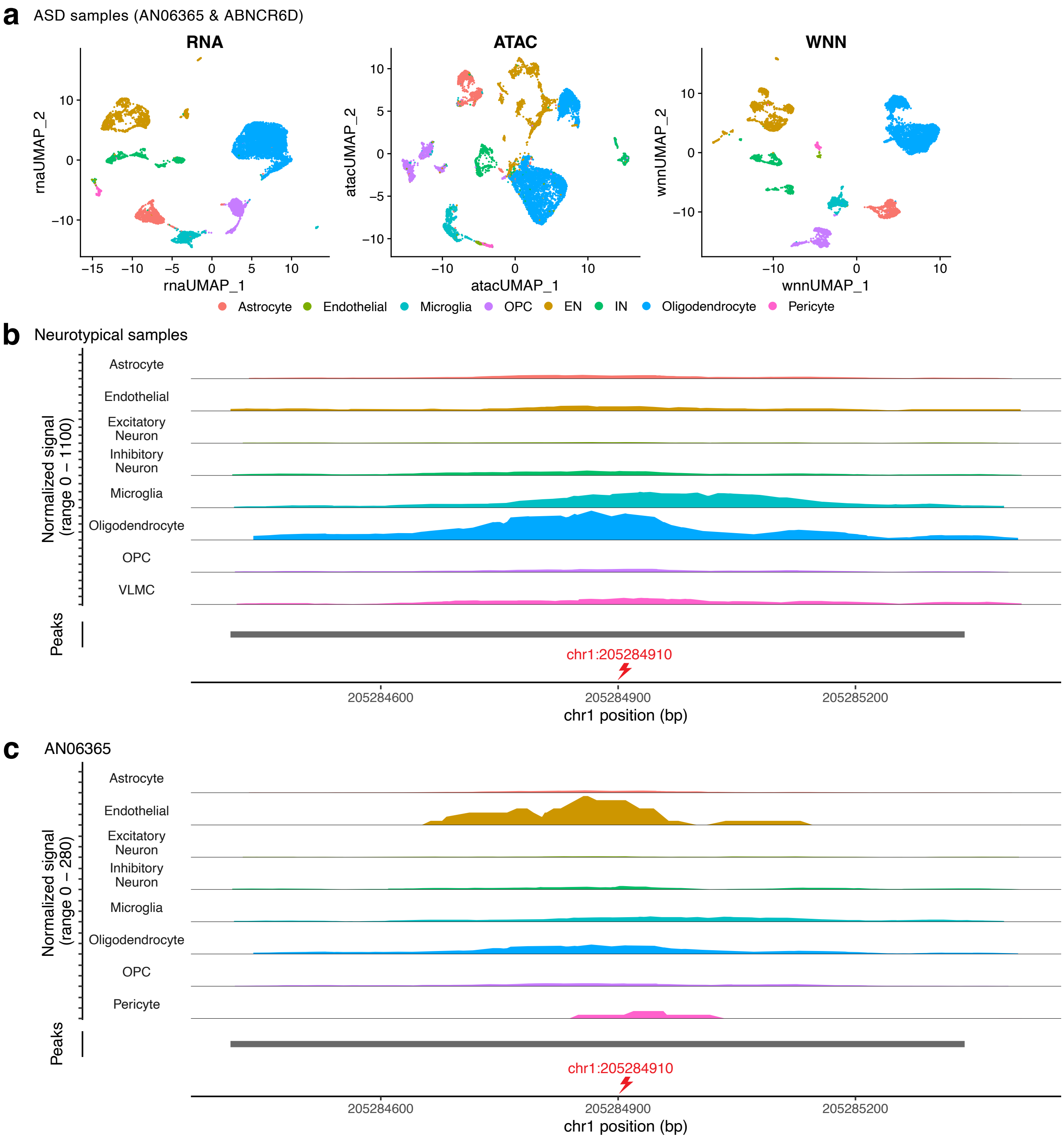


**Supplementary Fig. 3. Duplex-Multiome in ASD human brain.**

**Supplementary Fig. 3. Duplex-Multiome in ASD human brain.**

**a.** UMAP plots of snRNA-seq, snATAC-seq, and WNN-integrated data from two ASD brain samples. Cells are annotated and labelled by cell types based on their marker expressions.

**b.** Normalized ATAC signal across different cell types in neurotypical samples and AN06365. Peaks from snATAC-seq data are displayed.
